# RSV-induced Expanded Ciliated Cells Contribute to Bronchial Wall Thickening

**DOI:** 10.1101/2022.10.31.514471

**Authors:** Sattya N. Talukdar, Jaspreet Osan, Ken Ryan, Bryon Grove, Danielle Perley, Bony D. Kumar, Shirley Yang, Sydney Dallman, Lauren Hollingsworth, Kristina L. Bailey, Masfique Mehedi

**Affiliations:** Department of Biomedical Sciences, University of North Dakota School of Medicine & Health Sciences, Grand Forks, North Dakota, United States of America; Department of Internal Medicine, Pulmonary, Critical Care and Sleep and Allergy, University of Nebraska Medical Center, Omaha, Nebraska, United States of America

## Abstract

Viral infection, particularly respiratory syncytial virus (RSV), causes inflammation in the bronchiolar airways (bronchial wall thickening, also known as bronchiolitis), reducing airflow through the bronchioles. This bronchial wall thickening is a common pathophysiological feature in RSV infection, but it causes more fatalities in infants than in children and adults. However, the molecular mechanism of RSV-induced bronchial wall thickening remains unknown, particularly in healthy adults. RSV infection in the airway epithelium of healthy adult bronchial cells reveals RSV-infects primarily ciliated cells. RSV infection expands the cell cytoskeleton substantially without compromising epithelial membrane integrity and ciliary functions. The RSV-induced actin cytoskeleton expansion increases ununiformly epithelial height, and cytoskeletal (actin polymerization), immunological (INF-L1, TNF-α, IP10/CXCL10), and viral (NS2) factors are probably responsible. Interestingly, RSV-infected cell cytoskeleton’s expansion resembles a noncanonical inflammatory phenotype, which contributes to bronchial wall thickening, and is termed cytoskeletal inflammation.

**Author Summary:** RSV infects everyone. Although RSV-induced fatal pathophysiology (e.g., bronchiolitis) is more common in infants than adults, this bronchiolitis (or bronchial wall thickening) is common in the lower respiratory tract due to RSV infection in all ages. To determine the molecular mechanism of RSV-induced bronchial wall thickening, we infected *in vitro* adult airway epithelium with RSV. We found that RSV-infection induced a substantial actin-cytoskeleton expansion, consequently increased the height of the epithelium. We identified actin polymerization, secretion of proinflammatory cytokines and chemokines, and viral proteins contribute to the RSV-induced cytoskeletal expansion. Our results suggest that RSV-induces a novel noncanonical epithelial host response termed cytoskeletal inflammation, which may contribute to bronchial wall thickening.

## Introduction

Respiratory syncytial virus (RSV), a single-strand non-segmented negative-sense RNA virus, causes an enormous health burden on the healthcare system worldwide, as it increases morbidity and mortality in certain high-risk groups, including premature infants, infants with underlying medical conditions (such as chronic lung disease of prematurity and hemodynamically significant congenital heart disease), older adults, adults with chronic heart or lung disease, and adults with weakened immune systems [1–4]. Despite decades of research, no approved vaccines or cost-effective therapeutics against RSV are currently available.

RSV pathogenesis starts in the upper respiratory system via infection of airway epithelial cells (AECs) after RSV enters through the nostrils. The virus moves to the lower respiratory system and reaches the bronchioles, where viral replication induces pathophysiological mechanisms. RSV antigen is detected in both upper and lower respiratory track systems. In line with that, RSV is known to replicate in the epithelial cells of both upper and lower respiratory track systems [5–7]. RSV infection in the lower respiratory tract causes bronchial wall thickening in individuals of all ages [8–10]; however, the molecular mechanism of RSV-induced bronchial wall thickening, particularly in adults, remains unexplored. Bronchial wall thickening (bronchiolitis) is known as the inflammation of bronchioles [11]. However, the definition and diagnosis of bronchiolitis remain obscure due to a diverse group of etiologically, clinically, and pathologically dissimilar lesions [12]. Based on the microscopic pattern, bronchiolitis is diverse and can be broadly separated into acute and chronic categories [13]. While acute viral bronchiolitis may lead to severe bronchiolar epithelium necrosis [14, 15], common features of RSV bronchiolitis are edema, airflow obstruction, and mucus over-secretion [16]. Higher bronchial wall thickness is associated with multiple respiratory complications, including coughing and wheezing [17]. RSV bronchiolitis can increase the risk of developing asthma in children in their later stages of childhood [18]. In addition, RSV infection can be persistent and deteriorate lung function of COPD adults [19, 20]. Therefore, it is worth to elucidate the mechanism behind RSV-induced bronchial wall thickening.

RSV infection is responsible for virus-induced pathogenesis, which starts with virus entry. Two surface glycoproteins: glycoprotein (G) and fusion (F), play a significant role in RSV entry by host cell receptor binding and fusion, respectively [21, 22]. RSV structural proteins, including nucleoprotein (N), phosphoprotein (P), RNA-dependent RNA polymerase (L), and viral polymerase cofactor (M2-1), are involved in viral replication and transcription as well as its regulation, whereas M protein is mainly involved in viral assembly [23]. In contrast, RSV nonstructural proteins, including NS1 and NS2, are not the component of virion but play a significant role in viral pathogenesis by inhibiting Interferon (IFN)-I and III and inducing multiple cytokines and chemokines including RANTES, IL-8, and TNFα [24, 25]. The host actin cytoskeleton contributes to the different stages of RSV life cycles, including entry, transcription, replication, assembly, and budding [26]. RSV proteins interact with the host actin cytoskeleton and actin-associated or actin-regulatory proteins for its survival. RSV utilizes its ribonucleoprotein complex for replication in the host cytoplasm and a cytoplasmic structure named inclusion bodies formed by multiple viral proteins including N, P, L and M2-1 [27]. The obvious virus-induced modulation of ARP2/3-complex driven actin polymerization was evident in the filopodia-driven RSV cell-to-cell spread [28, 29]. However, RSV-induced cytoskeleton changes were evident in the in vitro 2D cell culture model. Thus, it is prudent to determine whether RSV-induced cytoskeleton modulation can be recapitulated in the more relevant 3D cell culture, e.g., air-liquid interface culture.

The aim of this study was to determine the molecular mechanism of RSV-induced bronchial wall thickening. Fort that, we used *in vitro* adult airway epithelium (3D cell culture), which resembles the epithelial airways *in vivo* with regard to morphology and functions, including mucus production, ciliary function, and membrane barrier integrity, but lacked immune cells (e.g., dendritic cells, macrophages, and T cells) [30–34]. We demonstrated that RSV infection expands the actin cytoskeleton and increases the airway epithelial cell layer height in the epithelium. RSV-infected epithelium shows resiliency, despite cytoskeletal expansion; in addition, RSV-induced epithelial height increases through a process termed cytoskeletal inflammation and indicate the existence of a noncanonical epithelial response to RSV infection. Our results suggest that RSV-induced cytoskeletal expansion is a novel mechanism of bronchial wall thickening.

## Results

### RSV infects ciliated cells, expands actin cytoskeleton, and increases height of the epithelium

To determine the mechanism of RSV-induced bronchial wall thickening in the adult airway epithelium *in vitro*, we first differentiated NHBE cells of three healthy adults independently for four weeks to form a well-differentiated pseudostratified mucociliary airway epithelium, which contained all three main cell types of the respiratory epithelium [32, 34, 35]: epithelial cells extending to the surface with cilia on the apical side, goblet cells containing mucinogen granules, and basal cells, which were confined to the basal portions of the epithelial layers on collagen-coated Transwells (Figs S1A&B). To confirm whether the passaging of NHBE cells up to four times did substantially change in the transcriptome profile, we collected total RNA from each passage cells and determined the whole-genome transcriptome profile by whole genome RNAseq analysis. We found no or less changes in the transcriptome profile passage to passage (Fig S1C); likewise, the cells retained normal epithelial phenotypic characteristics (Fig S1B) [32]. Additionally, the airway epithelium typically contains adherens, tight, and tricellular junctions [36, 37] that form an impermeable barrier; the presence of these junctions in our airway epithelium model was confirmed by detection of the epithelial intercellular junction protein E-cadherin, the peripheral membrane protein zonula occludins (ZO)-1, and the tricellular junction protein tricellulin (also known as MARVELD2), respectively (Fig S1D). We then infected this airway epithelium with RSV wild-type (RSV-WT) at a multiplicity of infection (MOI) of 4 for six days, which seems to be a standard duration for research on RSV infection in the airway epithelium [31, 38]. We compared the RSV-WT-infected airway epithelium with the uninfected (mock-infected) control epithelium under transmission electron microscopy [39] and found no evidence of gross cytopathic effects (CPEs) or culture deterioration (Fig S2A). However, compared to the mock-infected respiratory epithelium, the RSV-infected respiratory epithelium showed a few phenotypic differences, e.g., disorganization of granules in goblet cells, the appearance of higher interdigitation (cell membrane folding between cells), and an increased density of microvilli (Fig S2A). We confirmed RSV infects primarily ciliated cells by detecting RSV N mRNA in the infected ciliated but not in the goblet or basal cells (Fig S2B). We detected RSV virions on the apical sides of the infected ciliated cells, which suggests that RSV virions bud out from such cells. It appeared that RSV filamentous virion is too large to bud out from a cilium (Fig S2A, magnified). The possibility that RSV budding independent of cilium from the infected-ciliated cell is supported by a previous report of RSV budding from surface membrane microdomains in the nasal epithelium [40]. Therefore, we hypothesized that RSV shedding is independent of cilium in the bronchial epithelium. To confirm RSV shedding independent of a cilium from the infected ciliated cells, the RSV-WT-infected (MOI = 0.5) airway epithelium was fixed, permeabilized, and stained for the RSV fusion (F) protein and the ciliated cell marker acetyl-α-tubulin at 3 DPI. Indeed, we found that filamentous RSV virions budded out from the apical sides of the infected ciliated cells and independent of cilium (Fig S3A). No infectious RSV virions were detected in the basolateral medium by RSV immunoplaque assay, which suggests that virions are released from the apical surface of the airway epithelium; this finding is in line with those in previous reports [30, 31]. We also confirmed RSV primarily infected ciliated cells [30, 31] similar to human metapneumovirus (HMPV) [41] but in contrast to influenza A virus, which infected both ciliated and goblet cells [42] (Fig S2B, S3A-C). Expectedly, RSV’s unique cell tropism differed from that of different coronaviruses, e.g., severe acute respiratory syndrome coronavirus 2 (SARS-CoV-2) and Middle East respiratory syndrome (MERS), as both viruses infected preferentially goblet cells (Fig S3D) [35, 43, 44]. One of the striking differences we found was that RSV-infected cells were substantially enlarged, evident by the expanded actin cytoskeleton (Fig 1A and S3C). For better visualization of the RSV-induced expanded cytoskeleton, we infected the epithelium with recombinant RSV expressing green fluorescence protein (RSV-GFP) at an MOI of 4 for six days. We found that the RSV-GFP-induced expanded cytoskeleton was common to every infected cell (every GFP^+^ cell) in the epithelium model (Fig 1A and S4A and B). We then quantified the individual cell area and perimeter via Fiji [45] with the plugin PaCeQuant [46] (Fig 1B and C, S4B). RSV infection in the airway epithelium model increased the infected cell area and perimeter over time between 3 and 6 DPI (Fig 1B and C). The cell area and perimeter tended to increase over time in the infected cells within the infected epithelium.

**Fig 1.**
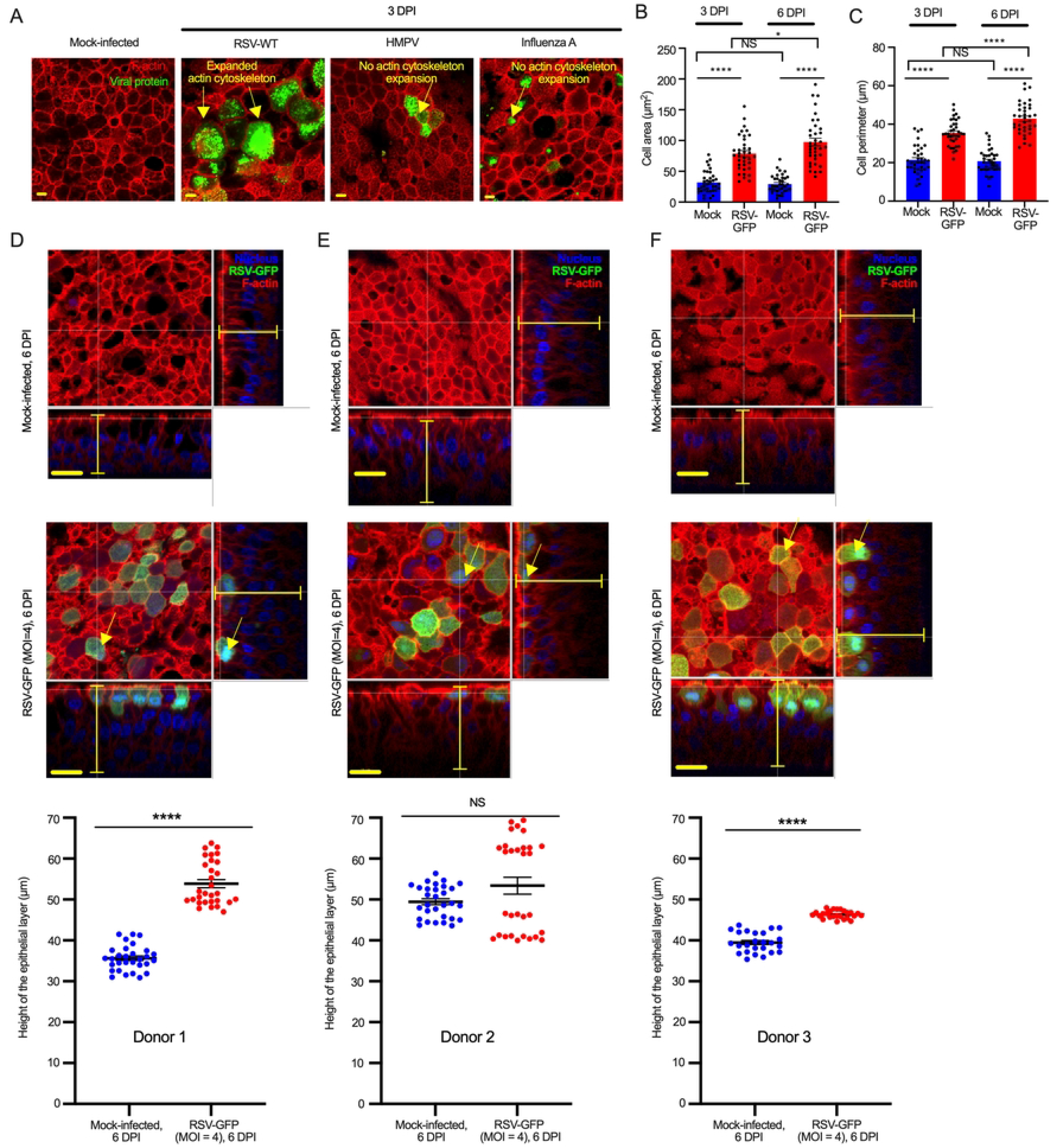
RSV-induced expanded-cytoskeleton increases bronchial wall thickening. (A) The airway epithelium was mock-infected or infected with different respiratory viruses (RSV-WT, HMPV, or influenza A virus PR8) independently at an MOI of 0.5 for 3 days. RSV F, HMPV N, or influenza A virus PR8 N were detected by immunofluorescence. F-actin (red) was detected by rhodamine phalloidin staining. The arrowhead indicates expanded actin cytoskeleton in the RSV-infected cell. The scale bar is 5 μm. (B-C) Cell areas and cell perimeters values were determined based on the assessment of rhodamine phalloidin in multiple random areas to cover most cells in the image (at least 36 random cells). The data represent at least two independent experiments. The error bars represent the SEMs. Statistical significance was determined by a two-tailed unpaired t-test. *, p<0.05; **, p<0.01, ***, p<0.001; ****, p<0.0001; NS, Not Significant. (D-F) Epithelium height was measured by confocal Z-stack imaging of mock- or RSV-GFP (MOI = 4)-infected epitheliums at 6 DPI. The data were obtained from 3 donors (Donor 1, 2 and 3). (D-F, Top and Middle) Representative images of mock- and RSV-GFP-infected (MOI = 4) epithelium at 6 DPI obtained from 3 donors. F-actin (red) and nuclei (blue) were stained with rhodamine phalloidin and DAPI, respectively. The scale bar is 15 μm. Yellow capped line and yellow arrow indicate the distance from basal to apical part of epithelium and RSV (RSV-GFP)-infected cell, respectively. (D-F, Bottom) To determine the epithelium height of Donor 1 and Donor 2, data was obtained from two independent experiments and the epithelium height of Donor 3 was determined from one independent experiment. The error bars represent the SEMs. Statistical significance was determined by a two-tailed unpaired t-test. *, p<0.05; **, p<0.01, ***, p<0.001; ****, p<0.0001; NS, Not Significant.

To determine the implications of the actin cytoskeleton expansion, we measured the thickness of the respiratory epithelium under confocal microscopy. We found that compared with mock infection, RSV infection significantly thickened the airway epithelium (three independent donors) (Fig 1D-F). The increment of epithelium height was observed higher at 6 DPI and the area where the infected cells showed higher increments (Fig 1D-F). No obvious increases in cell number were observed in the RSV-infected epithelia under confocal microscopy, suggesting that the observed thickening of the infected epithelia could have been due to enlargement of RSV-infected cells. Overall, the findings show that the RSV-induced increases in epithelial cell layer contribute to epithelial cell layer thickening, which may resemble bronchial wall thickening.

### RSV-infected thickened-airway-epithelium is resilient

To determine whether the presence of substantially enlarged infected cells impacted the biophysical properties of the respiratory epithelium, we first assessed the membrane integrity by evaluating the expression/abundance of three junction proteins (E-cadherin, ZO-1, and tricellulin) between the mock-infected and RSV-GFP-infected epithelia at 6 DPI. RSV infection did not reduce the expression of these proteins at 6 DPI (Fig 2A and S5A-C), indicating that membrane integrity was maintained. To assess whether the membrane permeability of the infected epithelia was stable throughout the infection process, we measured transepithelial electrical resistance (TEER) daily up to 6 DPI. TEER measurements, while very effective and powerful, are subject to variability. To obtain reliable impedance measurements, we subtracted the resistance measurement of a blank. We found that RSV infection did not reduce the membrane permeability of the infected epithelia relative to that of the mock-infected epithelia (Fig 2B). We then determined whether RSV infection disrupts ciliary motion at the apical surface of the epithelium. Any disruption of ciliary beating may indicate compromised ciliary function and respiratory disease progression. To quantify ciliary beat frequency (CBF), we used high-speed video microscopy with a Leica DMi8 epifluorescence microscope attached to an environmentally controlled chamber and analyzed the video files with Sisson-Ammons Video Analysis (SAVA) software V.2.1.15 (Ammons Engineering, MI, USA). We measured the CBF before and after infection of each Transwell without adding any medium to the apical side of the epithelium. We found that RSV-GFP infection in the airway epithelium did not reduce CBF over time (tested up to 6 DPI), except for a slight decrease at 6 DPI (Fig 2C). This reduction at 6 DPI probably due to deteriorating culture condition rather than viral increase over time, as RSV infection has been found to persist for longer than 6 days in a human airway epithelium model without obvious cytopathogenesis [30].

**Fig 2.**
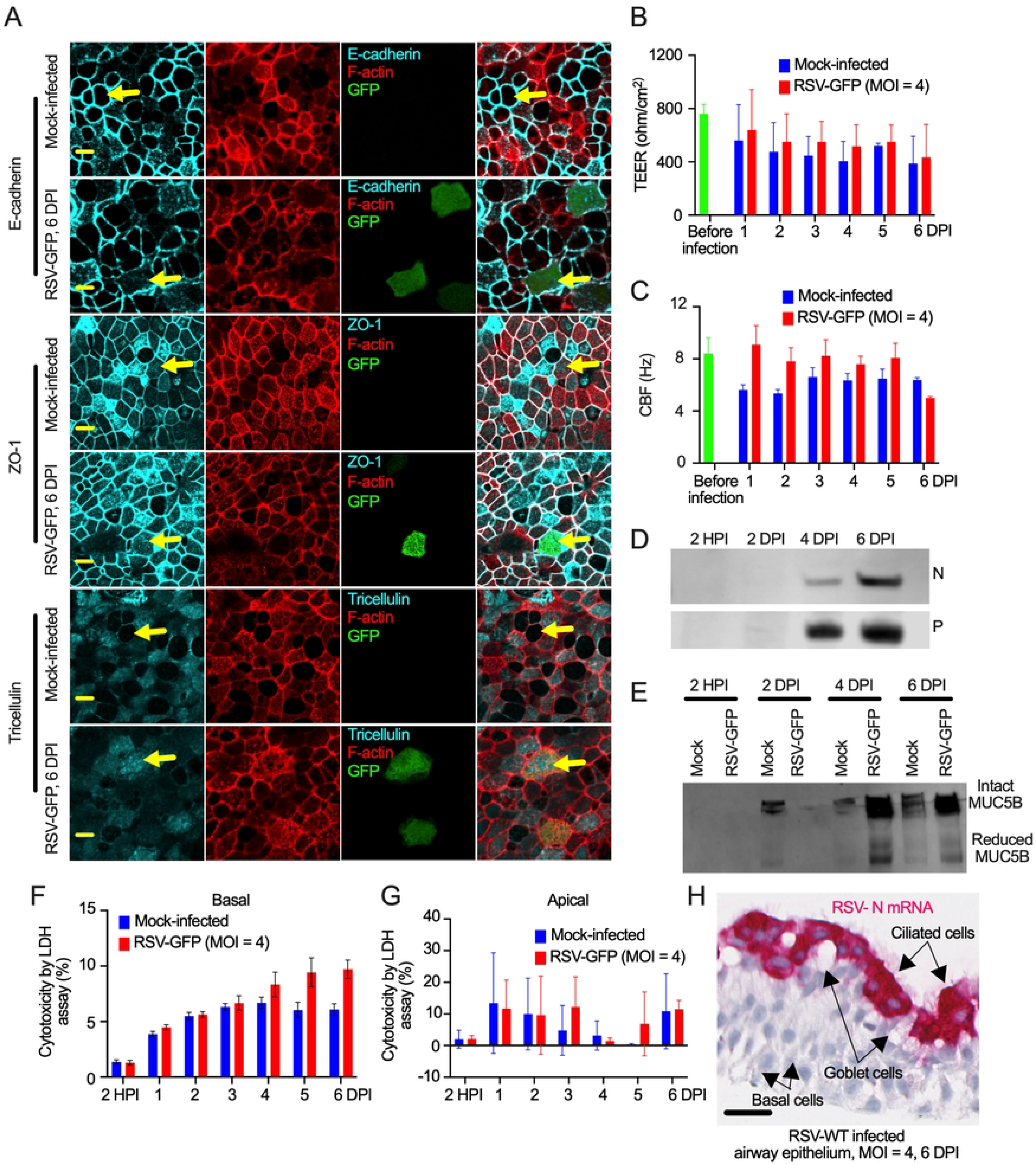
Resilience of the RSV-infected respiratory epithelium. (A) At 6 DPI, mock- and RSV-GFP-infected (MOI=4) epithelium were fixed, permeabilized, and stained with primary Abs against E-cadherin, ZO-1 and tricellulin followed by the respective secondary Abs. The images represent at least two independent experiments. The scale bar is 5 μm. Yellow arrow indicates the expression of E-cadherin, ZO-1 and tricellulin with and without RSV infection (B) The TEER values of mock- and RSV-infected epithelium were measured immediately before infection and at 1, 2, 3, 4, 5 and 6 DPI with a volt-ohm meter. The graph represents data combined from two independent experiments. For each experiment, the data were obtained by combining three independent Transwell reads, and each Transwell read was an average of three independent replicate reads. The green bar represent average read from twelve independent Transwells before infection. The error bars represent the SEMs. (C) The CBFs of mock- and RSV-infected epithelium was measured by high-speed video microscopy followed by quantification using a SAVA system before infection and at 1, 2, 3, 4, 5 and 6 DPI. The graph represents data combined from two independent experiments. For each experiment, the data were obtained by combining three independent Transwell reads, and each Transwell read was an average of reads from six random points. The green bar represent average read from twelve independent Transwells before infection. The error bars represent the SEMs. (D) Apical washes were collected at 2 HPI and at 2, 4, and 6 DPI from RSV-GFP-infected (MOI=4) epithelium, and RSV N and P protein levels were detected by Western blotting using specific Abs against N and P. (E) Mucin (MUC5B) production was detected by Western blotting (with reducing gels and heat treatment) of apical washes at 2 hours post infection (HPI) and at 2, 4, and 6 DPI for mock-infected or RSV-GFP-infected (MOI = 4) epithelium using Abs against MUC5B. (F and G) Basal medium (F) and apical washes (G) from mock- and RSV-GFP-infected (MOI = 4) epithelium was tested for cytotoxicity by LDH assay. (H) RSV N mRNA was detected using RNAscope 2.5 RED assay and anti-RSV N probe. Scale bar is 25 μm.

To determine whether RSV infection, particularly the presence of substantially enlarged infected cells impacts the biophysical properties of the respiratory epithelium, we assessed the overall viability of the infected epithelium by live-cell imaging from 5 DPI to 6 DPI. No apparent epithelial damage, cell sloughing, or substantial loss of ciliated cells was observed in the RSV-infected airway epithelium. Interestingly, a few RSV-infected (GFP+) cells disappeared from the infected epithelium, which indicates infection resolved or infected-cell damage (Movies S1 and 2). It appeared that RSV-infected cells can survive longer, but the cell-to-cell virus spread is an inefficient process in the respiratory epithelium (Movies S1 and 2). To confirm an increase RSV replication over time in the epithelium for an indication of productive infection, we collected apical wash from the RSV-GFP-infected epithelium, and 10 μg of total protein was subjected to detect RSV structural proteins, such as nucleoprotein (N) and phosphoprotein (P), by Western blotting. We detected both N and P from 4 DPI onward and found that the levels of both proteins were higher at 6 DPI than at 4 DPI (Fig 2D). The increase in viral protein level in the apical wash of the infected airway epithelium confirmed that RSV replicated in this airway epithelium. Elevated mucus secretion is a hallmark of RSV bronchiolitis [6] that can be recapitulated in *in vitro* airway models [31]. We therefore determined mucus production by measuring MUC5B protein levels in apical wash following infection by Western blotting. We found that mucus secretion increased over the course of RSV infection up to 6 DPI (Fig 2E). We then investigated whether RSV infection induces cytotoxicity in the airway epithelium by quantifying lactate dehydrogenase (LDH) from both apical wash and basal medium every day following infection. We found that LDH release into the basal medium during early infection for the infected samples was comparable to that for the mock-infected control samples, except for a slight increase later during the infection (Fig 2F). When we measured LDH release in the apical wash, the release was similar throughout the experimental period between the RSV-GFP-infected group and the mock-infected control group (Fig 2G). Based on the LDH release data, RSV did not induce substantial cytotoxicity in the infected respiratory epithelium. To confirm whether airway epithelium intact after RSV infection, we stained formalin-fixed paraffin embedded RSV-WT infected airway epithelium (at 6 dpi) for RSV N mRNA using a N specific mRNA probe. Indeed, we found that the RSV-infected ciliated cells were sustained without an obvious damage of the epithelium, which is in line with the previous results described above (Figs 2H, S2B). Overall, these results indicate that the adult airway epithelium shows substantial resilience in response to RSV infection.

### RSV modulates actin signaling pathways in the airway epithelium

To determine the molecular mechanisms mediating the expansion of the actin cytoskeleton in the thickened RSV-infected adult airway epithelium, we determined the global (genome-wide) transcriptional response of the RSV-WT-infected epithelia by RNA-seq analysis at 6 DPI. Briefly, we compared the resulting transcriptome profile with that of the mock-infected epithelia. First, we confirmed the validity of the transcriptome analysis by principal component analysis (PCA) of the transcriptome data, which showed that there was substantial variation between the mock-infected and RSV-infected transcriptome profiles, while there was much less variation within each group (Fig S6A). Second, we determined and ranked a list of differentially expressed genes. The differential gene expression analysis showed that approximately 3200 genes were upregulated and that a similar number of genes were downregulated in the RSV-infected epithelia compared to the mock-infected epithelia (Fig 3A and Table S1). Third, functional enrichment analyses were performed for the differentially expressed genes using clusterProfiler and the signaling pathway impact analysis (SPIA) package from Bioconductor. Gene enrichment analyses performed using STRING (a protein-protein interaction network database) against the Gene Ontology (GO) dataset for biological processes showed that most of the top 26 biological processes were significantly associated with cytoskeletal regulation except for three, which were I-kappaB kinase signaling that are known to be associated with nuclear factor kB (NF-kB)-driven inflammation (Fig 3B and Table S2) [47]. Fourth, we generated an enrichment map to identify functional modules. Enrichment maps generally organize enriched terms into networks with edges connecting overlapping gene sets, which tend to cluster together [48]. Based on the enrichment map, we found a major cluster linked to actin filament organization (Fig 3C). We further evaluated the RSV-induced modulation of actin cytoskeleton regulation in the respiratory epithelium. We found that several genes of the actin-regulatory network were substantially upregulated, contributing to the upregulation of actin polymerization (Fig 3D and S6B-C and Table S3). Thus, RSV preferentially modulates actin signaling pathways, particularly actin-related protein ARP2/3 complex driven actin polymerization, which is in line with previous reports [28, 49, 50].

**Fig 3.**
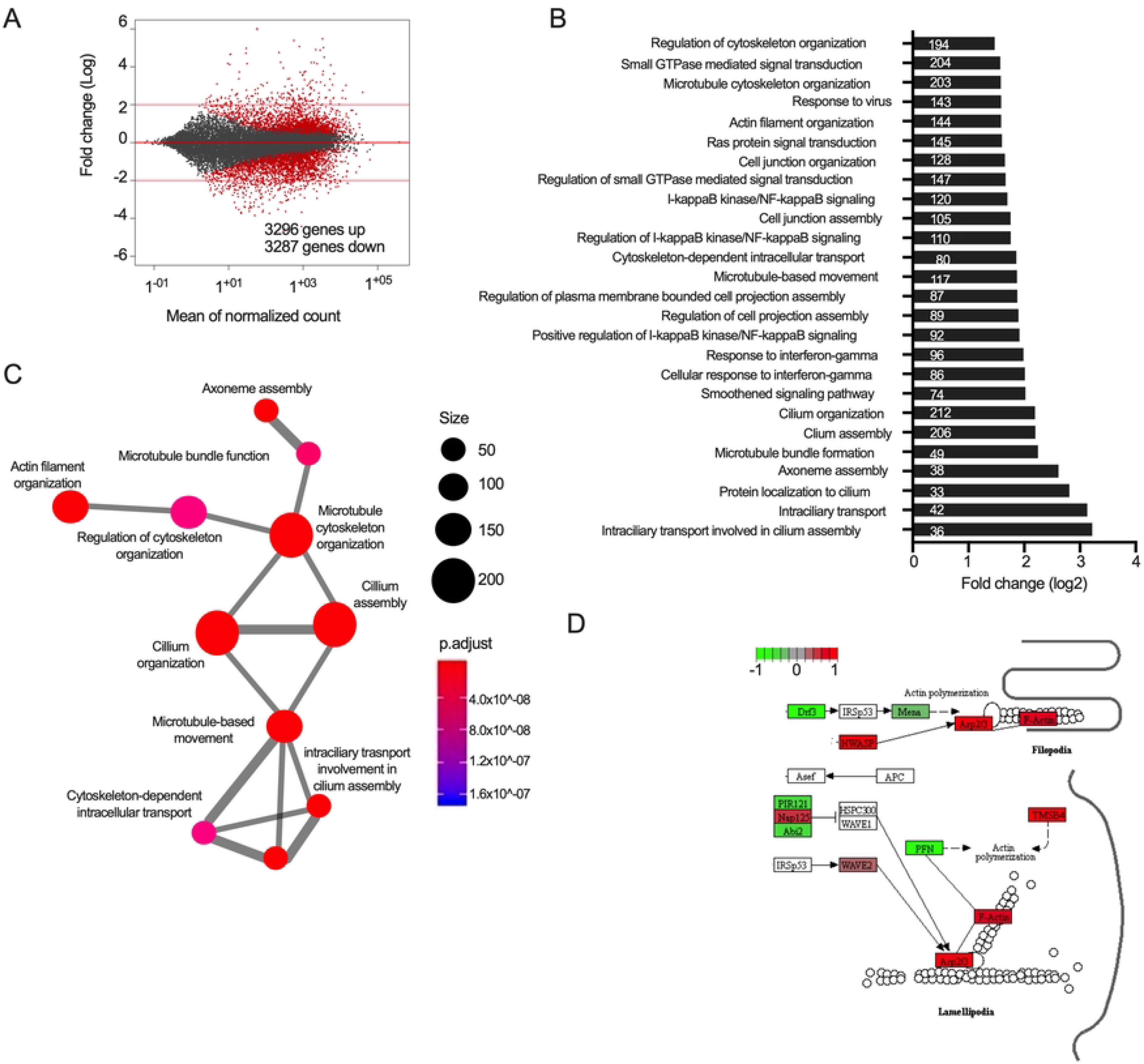
RSV modulates the actin-regulatory signaling pathway. (A) MA plot showing genes differentially expressed in RSV-WT-infected (MOI=4) epithelium compared with mock-infected epithelium at 6 DPI. Gray dots represent non-differentially expressed genes, and red dots represent differentially expressed genes. (B) Bar graph of the top 26 Gene Ontology (GO) biological process terms as determined by the R/Bioconductor package DESeq2 v1.24.0 [102]. The number of genes associated with each biological process is shown inside the bar. (C) Biological process interaction plot generated by creating an EMAP plot showing the interactions between different processes. The dot size represents the number of genes associated with a biological process. (D) SPIA enrichment analysis was used to evaluate the activation of the actin-regulatory pathway in RSV-infected samples. The Kyoto Encyclopedia of Genes and Genomics (KEGG) pathway of the partial actin-regulatory pathway is shown (see Fig S5B). Red represents upregulation, green represents downregulation, and white represents no change.

### ARP2/3 complex-driven actin polymerization contributes to RSV-induced cell expansion

The ARP2/3 complex is one of the three known actin nucleators in eukaryotes, which has a unique capability to organize filaments into branched actin network [51] for cell expansion [52] and it’s activity can be differentially regulated in cells [53]. RSV-driven modulation of ARP2/3 complex has already been shown in human lung epithelial A549 cells [28]. Here, we hypothesized that RSV-induced infected cell expansion depends on the ARP2/3 complex-driven actin polymerization. To determine RSV-induced enhanced actin polymerization, we quantified total F-actin / G-actin ratio between mock-infected and RSV-infected airway epithelium at 6 DPI. We found a substantial increase in F-actin levels compared to G-actin levels in the RSV-infected epithelium, and the F-actin/G-actin ratio was higher in the infected epithelium than in the mock-infected epithelium, indicating that higher F-actin levels were correlated with actin cytoskeleton expansion in the RSV-infected epithelium (Fig 4A and B). The increased actin polymerization mediated by ARP2/3 complex may corelate with the levels of F-actin, which plays a major role in organization of the apical actin cytoskeleton in polarized epithelium [54].

**Fig 4.**
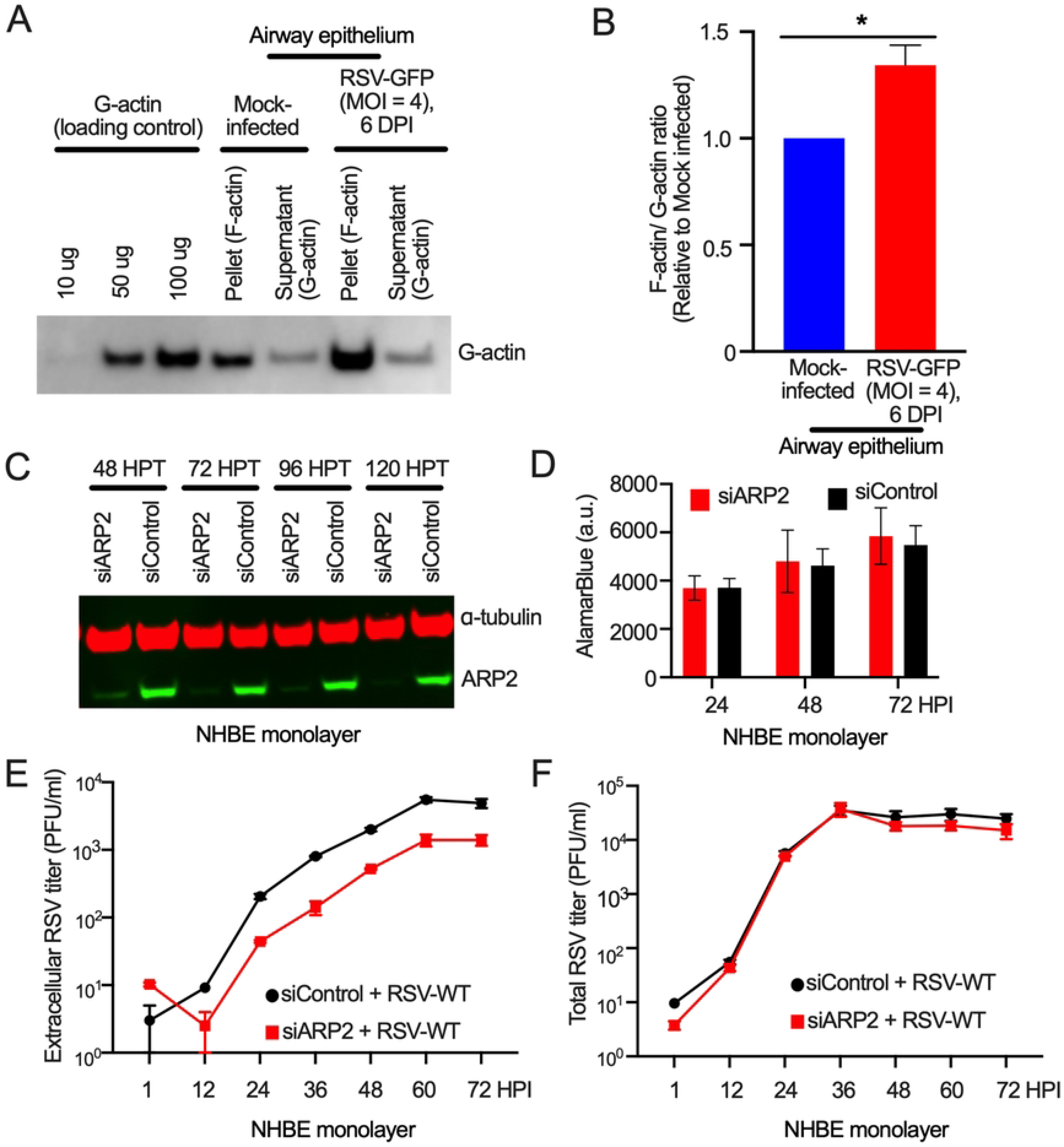
RSV-induced ARP2/3 complex-driven actin polymerization. (A and B) F-actin/ G-actin ratios was determined from the mock- or RSV-GFP infected (MOI = 4) airway epithelium at 6 DPI. (A) Various amounts of G-actin (10 ng, 50 ng, and 100 ng) were loaded for control. G-actin from the pellets (depolymerized F-actin) and supernatants of mock- and RSV-GFP-infected (MOI = 4) epithelium were loaded. G-actin was detected by Western blotting using a G-actin-specific Ab. (B) Bar chart shows the difference of F-actin/ G-actin ratio between mock- and RSV-GFP-infected (MOI = 4) 6DPI. The data were obtained from three independent experiments. The error bars represent the SEMs. Statistical significance was determined by a two-tailed unpaired t-test. *, p<0.05; **, p<0.01, ***, p<0.001; ****, p<0.0001; NS, Not Significant. (C) NHBE cell monolayer was transfected with either siARP2 or siControl for up to 120 hours. Western blotting for ARP2 detection was performed at 48-, 72-, 96- and 120-hours post transfection (HPT). (D) NHBE monolayer cells were transfected with either siARP2 or siControl for 72 hours. An alamarBlue assay was performed to determine cell viability at 24, 48, and 72 HPT. (E and F) NHBE monolayer cells were transfected with either siARP2 or siControl for 48 hours and then infected with RSV-WT (MOI = 0.5) for three days. The extracellular infectious RSV level was determined by titration of the supernatants from the infected cells. The total RSV level was determined by titration of both (E) the supernatants and (F) supernatants and cells togetherVirus titration was performed using an immunoplaque assay [28]. The data represent three independent experiments, each performed in triplicate. Titration was performed on each sample in duplicate. The error bars represent the SEM.

Among the actin-regulatory genes, *ACTR2* encodes actin-related protein 2 (ARP2), a major constituent of the ARP2/3 complex, which drives branched actin polymerization [51]. In addition to playing a physiological role in actin polymerization, ARP2 has also been shown to favor RSV spread *in vitro* by contributing to at least two processes: virus shedding and filopodia-driven cell-to-cell spread [28]. Therefore, we hypothesized that depleting ARP2 would reduce RSV shedding from infected cells. Due to a lack of success in depleting ARP2 in the airway epithelium model, we chose to deplete ARP2 in primary epithelial cells. We found that siRNA mediated ARP2 depletion in NHBE cells, which were the cells used to establish the respiratory epithelium model, was stable and nontoxic (Fig 4C and D). RSV-WT exerted substantial CPEs in the infected NHBE cell monolayer. We optimized an RSV-WT multi-step replicative cycle in NHBE cells by infecting cells with an MOI = 0.5 for three days. The NHBE cell monolayer was treated with either ARP2-targeting siRNA (siARP2) or control siRNA (siControl) for 48 hours and then subjected to mock infection or infection with RSV-WT (MOI = 0.5) for three days. Importantly, ARP2 depletion reduced RSV shedding from infected cells, as quantified by titration of extracellular RSV obtained from the supernatants of the infected NHBE cells over time (Fig 4E). However, the total viral titers for both supernatants and cells combined were not decreased by ARP2 depletion (Fig 4F). These results suggest that the contribution of ARP2 to RSV shedding in the NHBE cell monolayer, which was in line with the result in the previous report of RSV infected A549 cells [28].

### Robust induction of both cytokines and chemokines during RSV-induced cell expansion

To determine RSV-induced cytokines and chemokines, we collected both apical wash and basal medium separately from mock-infected or RSV-GFP-infected (MOI = 4) epithelium at 2-, 6-, 12- and 24-hours post infection (HPI) by repeated sample collections from the same Transwells. Then, we used a commercially available multiplex assay with a LegnedPlex Human Anti-Virus Response Panel (13 plex combination of cytokines and chemokines) (BioLegend) [55]. We detected RSV-induced secretion of multiple cytokines and chemokines in both apical wash and basal medium and the higher expression as observed at 24 HPI (Fig 5A and B). We repeatedly found IL-8 values for both mock and RSV-infection were way beyond the detection scale of the assay (shown in white, Fig 5A and B). We observed RSV-induced higher secretion of type III IFN lamda 1(IFN-L1), which is well known for its role in antiviral immune activity (Fig 5A and B) [56]. We found that IFN-L1 was induced as early as 6 HPI in the apical sup, where it took 24 HPI to detect in the basal medium (Fig 5C). The observed delay in the detection of IFN-L1 in the basal medium might have correlated with the time required for viral protein production. The apparent low levels of cytokine detection by the assay were more likely due to successive samples (described above) were collected from the same Transwell than to limitations of detection by the kit.

**Fig 5.**
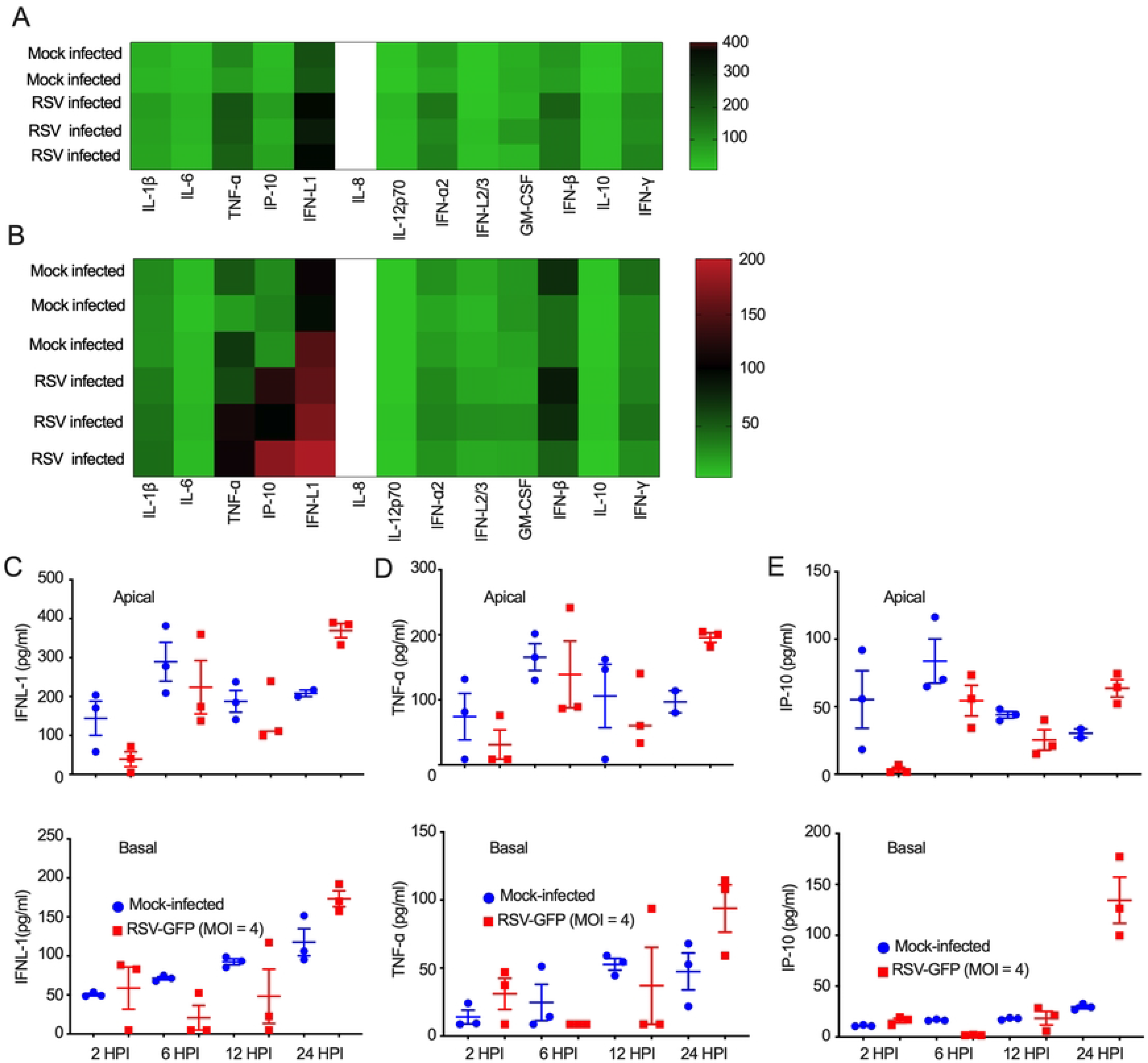
RSV induces substantial secretion of cytokines and chemokine. A LegendPlex Human Anti-Virus Response Panel was used to analyze both the apical and basal parts of mock- and RSV-GFP-infected (MOI=4) epithelium at early times post infection. The levels of cytokines and chemokines were determined in the apical supernatant (A) and basal medium (B) at 1 DPI. The data represent one independent experiments, each performed in triplicate. The levels of two cytokines IFN lamda-1 (C) and TNF-α (D) and one chemokine IP10/CXCL10 (E) were quantified at early times post infection. The data represent two independent experiments, each performed in triplicate. The error bars represent the SEM.

Along with cytoskeletal regulatory signaling, I-kappaB kinase signaling was one of the major biological processes modulated by RSV infection in the airway epithelium model (Fig 3B). The canonical activation of NF-kB depends primarily on I-kappaB kinase signaling [57]. To confirm the involvement of NF-kB signaling, we further analyzed our transcriptome data for gene enrichment via Kyoto Encyclopedia of Genes and Genomes (KEGG) pathway analysis and found that the tumor necrosis factor (TNF) signaling pathway was one of the top pathways modulated by RSV infection (Table S4). Activation of NF-kB by TNF-family cytokines in viral infection is well established [47, 58]. We found that several genes of the TNF signaling network were upregulated in our transcriptome data (Fig S7 and Table S5). Indeed, we found that TNF-α secretion was higher in both the apical wash and basal medium from the RSV-infected epithelium than in those from the mock-infected control epithelium at the early time point (Fig 5D). However, the detection of TNF-α in both apical wash and basal medium from the mock-infected control epithelium indicated the presence of nonspecific or inherent inductions.

NF-kB activation induces not only proinflammatory cytokines, e.g., TNF-α, but also proinflammatory chemokines, e.g., IP-10 (CXCL10) (Fig S7) [57]. We hypothesized that proinflammatory chemokines also contributed to the observed thickening of the RSV-infected airway epithelium. Indeed, we found substantial and robust IP-10 induction in the infected epithelium at the early time point of infection (Fig 5E). This robust IP-10 induction was also evident when quantified using a different multiplex assay with a LegendPlex Human Proinflammatory Chemokine Panel (BioLegend). The observed delay in the detection of IP-10 in the basal medium might have correlated with the time required for viral protein production/replication-dependent modulation of IP-10 signaling (Fig 5E, bottom). Importantly, the RNA-seq data indicated that the RSV-induced IP-10 secretion was sustained at the later time of infection (Table S1). These results suggest that RSV infection induced both proinflammatory cytokines (IFN-L1 and TNF-α) and chemokine (IP-10), whether these cytokines and chemokine contribute to the RSV-induced modulation of actin polymerization for the observed cell expansion remains to be determined.

### Robust expression of NS2 during RSV-induced cell expansion

To determine the contributions of viral proteins to the thickening of the RSV-infected airway epithelium, we analyzed RNA-seq data to assess all RSV genes for differential expression, which are available through the GEO under the accession number GSE146795. To quantify RSV gene specific transcripts, the reads from RNA-seq were aligned to the human RSV genome (GenBank accession KT992094) with STAR v2.7.1a [59]. We found that most of the genes encoding for structural and nonstructural proteins were substantially upregulated except M2-2 and L (Fig 6A). Importantly, NS2 transcript was the highest among all the viral transcripts. RSV nonstructural protein 2 (NS2) has already been implicated for RSV-induced pathophysiology [38]. Thus, we determined NS2 secretion from the RSV-infected airway epithelium. NS2 was detected in the apical supernatants as early as 2 days post infection and secretion were increased over time (Fig 6B and C). The lower detection of NS2 at 6 DPI than 4 DPI was probably as a result of repeated sample collection at different time points on the apical side of the same transwell. In addition to NS2, we also found elevated transcripts for multiple viral structural proteins: N, P, matrix (M), and F. We also detected and quantified a few of those proteins by Western blotting. Indeed, viral protein level increased over time, e.g., N and P levels were higher at 6 DPI than at 3 DPI (Fig S8A and B). As intracellular protein increased over time, the NS2 low detection at 6DPI explains due to the repeat sample collection. Overall, these data suggest that a number of RSV proteins, importantly NS2 expressed substantially during RSV-infected cell expansion. However, how NS2 directly contribute to RSV-induced modulation of actin polymerization for the observed cell expansion is remains to be determined.

**Fig 6.**
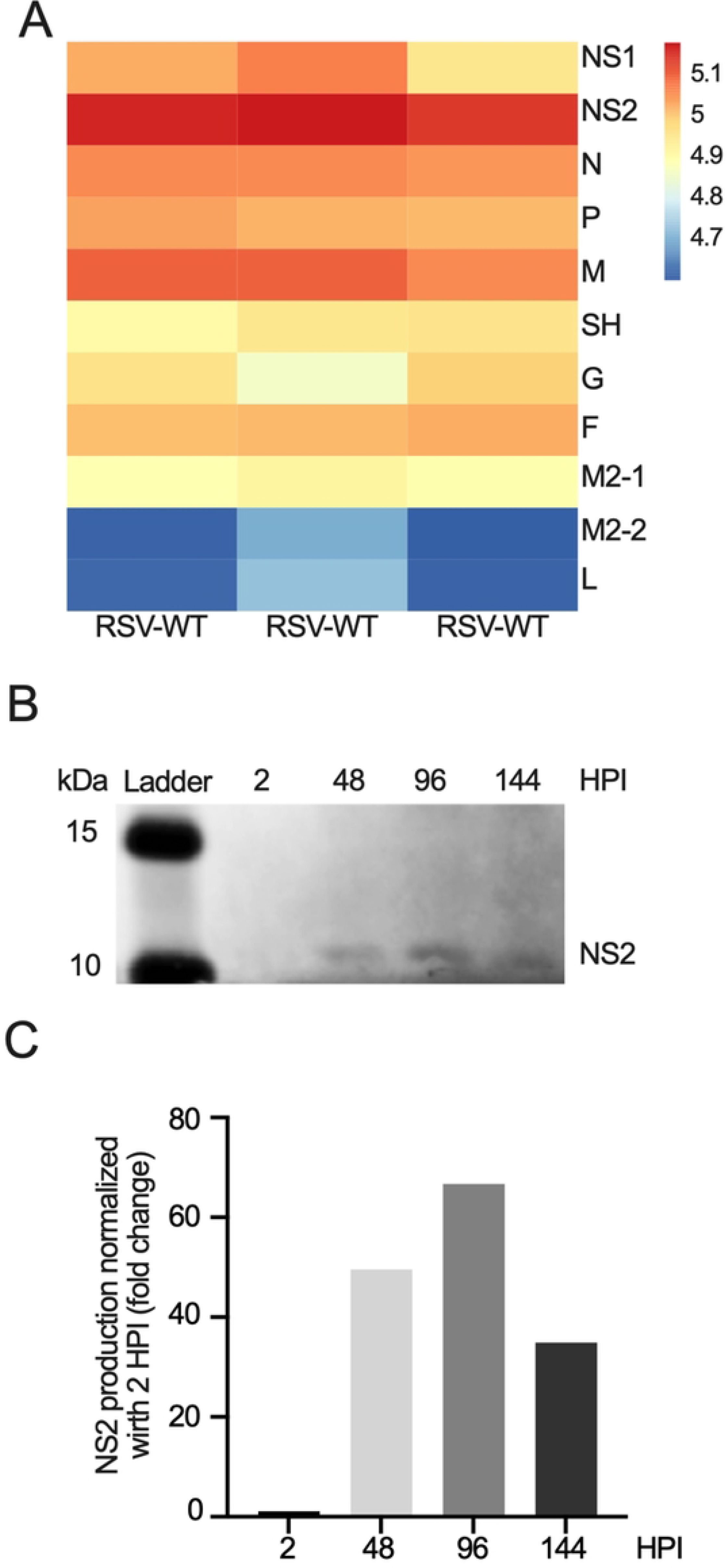
Robust NS2 secretion in the infected respiratory epithelium. (A) Airway epithelia were mock-infected or infected with RSV-WT (MOI = 4) for six days. RSV transcript the reads (RSA-seq) were aligned to the human RSV genome (GenBank: KT 992094). (B) Nonstructural protein 2 (NS2) production was detected by Western blotting and samples were collected from apical washes at 2 HPI and at 2, 4, and 6 DPI for RSV-GFP-infected (MOI = 4) epithelium using Abs against NS2. (C) NS2 protein production over time normalized to 2 HPI.

## Discussion

Here, we first established airway epithelium of three healthy adults NHBE cells independently with membrane integrity and ciliary functions, which are in line with previous publications [32, 34]. Previous studies have shown that both early-passage and later-passage primary NHBE cells from different donors can form airway epithelium with similar phenotypic characteristics [30–32]. In this study, we confirmed that passaging NHBE cells up to four times did not substantially change the whole-genome transcriptome profile (Fig S1C); likewise, the cells retained normal epithelial phenotypic characteristics [32]. Additionally, we found that neither passaging NHBE cells nor differentiating them into an airway epithelium required the presence of a Rock inhibitor or feeder cells [32, 60]. In separate studies, we also showed establishing an *in vitro* airway epithelium by using P4 NHBE cells of a chronic obstructive lung disease (COPD) patient without using of a Rock inhibitor or feeder cells [35, 61] .

We confirmed RSV infects primary ciliated cells in the adult airway epithelium without any obvious cytopathogenicity, which is in line with previous study [30]. We found that filamentous RSV virion budding was independent of cilium on the surfaces of the infected ciliated cells, in agreement with the observations of other researchers [62]. The most striking difference we found was that RSV infection, in contrast to HMPV or influenza virus A infection, expanded the actin cytoskeletons of the infected cells. While virus-induced modulation of actin has been well established *in vitro* [63–65], a functional cytoskeletal system may be dispensable for viral replication [66]. Specifically, the involvement of actin in RSV entry [67], replication [68], and spread [28] *in vitro* has already been established. RSV-induced changes in the infected cell morphology were shown in the human airway culture of tracheobronchial specimen from a lung transplant patient [38]. Here, we show that RSV infection in adult airway epithelium expands the actin cytoskeleton, which consequently increases the epithelial height and may provide a novel explanation of the mechanism of bronchial wall thickening *in vivo*. Bronchial wall thickening (airway inflammation) is a common finding in lower respiratory tract infections caused by RSV in infants, children, and adults [8, 9]. RSV-induced epithelial height increment is not always uniform, and it is probably due to a random spatial distribution of the infected ciliated cells and the degree of infection. As the human airway contains a higher percentage of ciliated cells than other types of cells, e.g., goblet cells, RSV-infection-induced cytoskeletal expansion and its contribution to bronchial wall thickening are primarily related to the degree and persistent of infection. Further research is necessary to determine how RSV infection’s spatial and temporal distribution in the bronchiolar airway epithelium contributes to bronchiolitis. SARS-CoV-2 infection apparently does not cause bronchiolitis [69]. Whether disparity in cell tropism cause different pathophysiology between SARS-CoV-2 and RSV infections yet to be determined.

Increases in cell size and number along with other factors, e.g., recruitment and activation of immune cells, have been shown to be contributing factors for bronchial wall thickening in chronic lung disease, such as asthma [70]. Viral infection is known to increase airway inflammation (in acute exacerbation or chronic inflammation), and immune cells are key players in this process. Here, we showed that how epithelial cells particularly RSV-infected ciliated cells contributed to the height increase of the airway epithelium and suggesting an existence of a noncanonical inflammatory phenotype in the airway epithelium. This noncanonical inflammatory phenotype can be explained by RSV-induced epithelial inflammatory phenotype by virus-infected ciliated cells without presence of any classical immune cells (monocyte, dendritic cells and macrophages). Thus, the inflammatory phenotype driven by RSV-induced cytoskeletal expansion without the presence of classical immune cells can be termed cytoskeletal inflammation, which may contribute to bronchial wall thickening – a pathophysiological feature of RSV infection in the lungs [8]. Thus, RSV-induced cytoskeletal inflammation may shed light on the mechanism of bronchiolitis in infants with immature immune responses [71].

The expanded actin cytoskeleton in the RSV-infected airway epithelium model showed strong resilience, as increased viral replication over time did not impact membrane integrity, ciliary functions, or cell sloughing. Although these fatal RSV pathophysiological features can be recapitulated in the pediatric bronchial epithelium [31], they were not recapitulated in the adult airway epithelium, except for elevated mucus secretion. MUC5B expression was significantly higher during RSV infection in this study which also observed in different in vivo and in vitro studies [72–74]. We believe that this discrepancy could be due to differences between the pediatric and adult airway epithelium, and it suggests that the airway epithelium in adults combats RSV infection better than those in infants and children. Alternatively, the virus strain, the MOI of the virus inoculum, or the duration of infection (up to six days) may have contributed to the observed discrepancy. However, RSV infection in the airway epithelium can be sustained for longer than six days without any obvious CPEs [30].

We identified potential cytoskeletal, immunological, and viral factors contributing to the cytoskeletal inflammation in response to RSV-induced bronchiolitis. With regard to cytoskeletal factors, we have shown that RSV modulates numerous genes relevant to cytoskeletal signaling, particularly ARP2/3, a complex of which drives actin polymerization. The higher F-actin in the RSV infected epithelium compared mock infection confirmed RSV-induced modulation of ARP2/3 complex dependent actin cytoskeleton signaling. RSV-induced modulation of ARP2/3 causes infected cell expansion in the airway epithelium, which is different from RSV-induced filopodia induction in A549 cells [28]. Neither syncytia formation nor cell-to-cell spreads was evident in the RSV infected airway epithelium, but both are common in the RSV infection in A549 cells [28]. Although it is obvious that RSV modulates ARP2/3 complex-driven actin cytoskeleton signaling in both 2D (A549 cells) and 3D (airway epithelium) culture models, the observed phenotypical differences are probably due to the generic difference between the models. No or less syncytia in the RSV-infected airway epithelium resembles clinical feature of RSV infection [75]. Importantly, ARP2/3 complex driven actin polymerization requires in RSV replication has been evident both lung epithelial A549 cells and primary NHBE cells. Thus, our result provides further evidence in support of the role of ARP2 in RSV replication cycle [28]. With regard to immunological factors, we found that RSV infection induces robust secretion of both cytokines (particularly, IFN-L1 and TNF-α) and chemokine (particularly, IP10/CXCL10). RSV-induced higher IFN-L1 level was also observed in previous studies [76, 77]. We also found a substantial induction of proinflammatory cytokine TNF-α, which is part of the NF-kB signaling pathway and is considered the central activator of inflammation and innate immunity in response to RSV infection [57, 78, 79]. In our studies, the most reliable and robust chemokine induction was IP10/CXCL10 and IL8 (above the saturation of detection limit), which we quantified by using two independent commercially available panels: a Human Anti-Virus Response Panel and a Human Proinflammatory Chemokine Panel. The results were consistent with previous reports of both *in vitro* and clinical findings [31, 80, 81]. In addition, multiple studies reported IP10/CXCL10 and IL8 are the most abundant chemokines caused by RSV infection [82, 83]. The elevated level of IP10/CXCL10 and its receptor CXCR3 has shown increasing leukocyte recruitment in different respiratory complications including COPD and peribronchiolar inflammation which eventually leads to airway fibrosis [84, 85]. The IP10/CXCL10-CXCR3 axis has regulatory role on respiratory epithelium by maintaining chemotaxis mediated by a Ca^2+^-independent and G-protein dependent pathway by inducing PI3K and p38 MAPK signaling pathways in human airway epithelial cells [86]. TNFα-induced elevated IP10/CXCL10 secretion also observed in COPD which is NF-kB dependent [87]. With regard to viral factors, we found that NS2 is substantially expressed in the RSV-infected epithelium, which is in line with a previous report of NS2 contribution to cell rounding phenotype [38]. Nonstructural protein-mediated actin rearrangement during RSV infection is not uncommon, rather several other bacterial and viral non-structural proteins demonstrated their interaction with actin cytoskeleton and their stimulating role for host cellular actin remodeling [88–93]. RSV-infection induces actin polymerization for increasing epithelial layer as we hypothesized whereas most of the abovementioned viruses have negative impact on actin polymerization for their growth. Our result demonstrated RSV infection stimulates higher TNFα secretion and higher F-actin/G-actin ratio indicating substantial actin polymerization which is identical with the previous study showing the induction of TNFα during airway inflammation was associated with the increase of total actin concentration and F-actin/G-actin ratio [94]. NS2-deleted recombinant RSV infection showed reduced NF-κB activation and lower TNFα production compared to wild-type RSV infection [25]. The possibility of NS2’s contribution to increase actin polymerization can be predicted because Listeria ActA protein driven ARP2/3 complex activation has already been characterized [93]. NS2’s role in the RSV-induced ARP2/3-complex-driven actin polymerization remains to be determined. Additionally, we found that RSV structural proteins, particularly N, P, M, and F, are essential for viral replication and are highly expressed. M- or F-induced F-actin interactions or modulation, respectively can be established *in vitro*, suggesting that these proteins may play a role in RSV-induced cytoskeletal expansion [28, 95].

In conclusion, we found that RSV infects primarily ciliated cells in the adult airway epithelium and showed a noncanonical inflammatory phenotype which resembles bronchial wall thickening. This RSV-induced cytoskeletal inflammation is due to upregulation of ARP2/3 complex driven actin polymerization, and robust induction of proinflammatory cytokines (IFNl-1 and TNF-α) and chemokine (IP10/CXCL10), and robust secretion of viral proteins, particularly NS2. The thickened airway epithelium shows resilience, which supports the existence of a noncanonical mechanism of bronchial wall thickening due to RSV infections in adults.

## Materials and Methods

### Primary cells, cell line, and virus

Primary NHBE cells (from 3 healthy nonsmoker adult donors) (deidentified) were provided by Dr. Kristina Bailey, at the University of Nebraska Medical Center (UNMC), Omaha, NE under an approved material transfer agreement (MTA). Vero cells (an African green monkey kidney epithelial cell line, ATCC-1586) and the viruses RSV-WT (A2 strain, GeneBank ID 2992094) and RSV-GFP (in which the GFP gene was inserted between the P and M genes of RSV-WT) were obtained from Dr. Peter Collins at NIH. Both viruses were grown separately in Vero cells and sucrose-purified using density-gradient ultracentrifugation. Virus stocks were stored at −80°C and tittered by immunoplaque assay using a cocktail of three monoclonal Abs specific to RSV F [28, 96]. HMPV was obtained from Dr. Peter Collins at NIH. The influenza A virus PR8 strain was obtained from Dr. Nadeem Khan at UND.

### Primary cells subculture

NHBE cell monolayer subculturing was performed in a 100-mm culture dish (Corning, Inc.). Briefly, a culture dish was coated with PureCol (Advanced Biometrics) before seeding of cryopreserved (−80°C) passage-one (P1) NHBE cells (approximately 10^6^ cells), which were thawed in a 37°C water bath. The cells were maintained in complete AEC growth medium (PromoCell) containing AEC supplement (PromoCell), 2% penicillin/streptomycin (Thermo Fisher Scientific) and 1% amphotericin B (Thermo Fisher Scientific) at 37°C in a 5% CO_2_ incubator. The cells were grown to 90% confluency, with medium changes every other day. Confluent monolayers of cells were dissociated with TrypLE (Thermo Fisher Scientific) for 5 min at 37°C and pelleted, and one-third of the cells were reseeded in a culture dish containing AEC medium with supplements for subculturing.

### Air-liquid interface culture

Transwells (6.5 mm) with 0.4-μm-pore polyester membrane inserts (Corning Inc.) were coated with PureCol for 20 min before cell seeding. NHBE cells (5×10^4^) suspended in 200 μl of complete AEC medium were seeded in the apical part of each Transwell. Then, 500 μl of complete AEC medium was added to the basal part of the Transwell. When the cells formed a confluent layer on the Transwell insert, the AEC medium was removed from the apical insert, and PneumaCult-ALI basal medium (Stemcell Technologies) with the required supplements (Stemcell Technologies), 2% penicillin/streptomycin and 1% amphotericin B was added to the basal chamber. The ALI medium in the basal chamber was changed every other day. The apical surface was washed with 1x Dulbecco’s phosphate-buffered saline (DPBS) (Thermo Fisher Scientific) once per week initially but more frequently when more mucus was observed on later days (when it was difficult to see the apical cells, it was determined that the difficulty was probably due to a thick layer of mucus). All cells were differentiated for up to four weeks (at 37°C with 5% CO_2_) until the desired cellular and physiological properties of an epithelial layer were obtained, such as a CBF greater than 6 Hz and a TEER greater than 500 ohm/cm^2^.

### Virus infection

The four-week-cultured highly differentiated pseudostratified airway epithelia were washed with 200 μl of 1x DPBS to remove mucus and infected on the apical side with sucrose-purified RSV-WT, HMPV, or influenza virus PR8 strain at an MOI of 0.5 in 1x DPBS for two hours (at 37°C with 5% CO_2_). The viral inoculum was then removed, and the epithelia were washed twice with 200 μl of 1x DPBS. Fresh ALI medium (500 μl) with supplements was added to the basal part of each Transwell, and the apical part was kept empty. Mock-infected (1X DPBS without virus) and RSV-infected Transwells were incubated for three days at 37°C in an incubator. A similar approach was used to infect the epithelia with RSV-GFP or RSV-WT at an MOI of 4 for six days. SARS-CoV-2 or MERS infection in the airway epithelium was described previously [35, 97].

### Confocal microscopy

The airway epithelium (apical side) was washed with 1xPBS, and both the apical and basal parts were fixed with 4% paraformaldehyde (PFA) (Polysciences, Inc.) in 1x PBS for 30 min at room temperature (RT). The epithelium was then washed twice with 1x PBS and blocked with a 10% goat serum solution in immunofluorescence (IF) washing buffer (130 mM NaCl_2_, 7 mM Na_2_HPO_4_, 3.5 mM NaH_2_PO_4_, 7.7 mM NaN_3_, 0.1% BSA, 0.2% Triton-X 100 and 0.05% Tween-20) for 1 hour at RT. After 2 washes with 1xPBS, the Transwell inserts were incubated with one of the following primary Abs in IF wash buffer overnight at 4°C: anti-ZO-1 rabbit polyclonal (1:200) (40-2200, Thermo Fisher Scientific), anti-E-cadherin rabbit monoclonal (1:200) (3195S, Cell Signaling Technologies), anti-MUC5B rabbit monoclonal (1:500) (HPA008246, Atlas Antibodies), anti-MARVELD2/ Tricellulin rabbit polyclonal (1:100) (48-8400, Thermo Fisher Scientific), anti-acetyl-α-tubulin rabbit monoclonal (1:500) (5335, Cell Signaling Technologies), anti-MUC5AC mouse polyclonal (1:100) (H00004586-A01, Abnova), anti-Respiratory Syncytial Virus F mouse monoclonal (1:100) (ab43812, Abcam), anti-Metapneumovirus N mouse monoclonal (1:25) (MCA4674, Bio-Rad), or anti-Influenza A nucleoprotein (NP) mouse monoclonal (1:30) (MCA400, Bio-Rad). The next day, after washing with the wash buffer, the inserts were incubated with an anti-rabbit Alexa Fluor 647-conjugated secondary Ab (1:200) (Thermo Fisher Scientific) and anti-mouse Alexa Flour 488-conjugated secondary Ab (1:200) (Thermo Fisher Scientific) in IF wash buffer for 3 hours in the dark at 4°C. The cells were washed 2 times with 1xPBS and then incubated with rhodamine phalloidin (PHDR1; 1:500) (Cytoskeleton Inc.) in IF wash buffer for 30 min at RT in the dark. After 2 washes with 1xPBS, the cell nuclei were stained with NucBlue Fixed Cell Stain ReadyProbes Reagent (4’,6-diamidino-2-phenylindole, DAPI) (Thermo Fisher Scientific) for 30 min in the dark at RT. The Transwell inserts were washed with 1xPBS, and then the entire membrane was removed from each well and placed on a microscope slide (TechMed services). ProLong Gold antifade reagent (Thermo Fisher Scientific) was used to mount coverslips on the slides. Images were captured using a confocal laser scanning microscope (Olympus FV3000) with a 60x objective. The 405-nm laser was used to excite DAPI for nucleus detection; the 488-nm laser was used to excite Alexa Fluor 488 (RSV Fstaining) or GFP for RSV detection; the 561-nm laser was used to excite rhodamine phalloidin for F-actin detection; and the 640-nm laser was used to excite Alexa Fluor 647 for MUC5B, acetyl-α-tubulin, E-cadherin, ZO-1, or tricellulin detection. At least three independent fields were chosen for imaging from each experiment. Slides prepared from more than two independent experiments were imaged. Imaris software version 9.5.1 (Oxford Instruments Group) was used for the conversion of Z-stack images (.oir format) to .tiff format and for additional image postprocessing.

### Paraformaldehyde (PFA)-fixation, paraffin embedding, and RNAScope

The airway epithelium (apical side) was washed with 1xPBS, and both the apical and basal parts were fixed with 4% paraformaldehyde (PFA). The PFA-fixed airway epithelium was then paraffin-embedded and section at 5 μm [35, 61]. 5μM section slides were first deparaffinized by incubating them following solutions in a coplin jar: 1. Histo-Clear for 5 min, x2; 2. 100% ethanol for 5 min, x3; 3. 95% ethanol for 5 min, x1; 4. 70% ethanol for 5 min, x1; and 5. distilled water for 5 min, until the next step. The deparaffinized slides were immediately incubated in 0.5% TritonX-100 solution in 1x PBS for 30 min in a coplin jar. The slides were washed three times with 1X PBST (1x PBS with Tween 20) for 5 min. A hydrophobic barrier was drawn around the 5μm section on the slides by using an Immedge Hydrophobic Barrier Pen. To reduce nonspecific antibody binding, the section was blocked 10% goat serum solution (Vector Laboratories) in 1x PBST for 2 hours at room temperature. The slides were then incubated with anti-RSV N mRNA probe (V-RSV-NP-01) and RNAscope 2.5HD Detection kit (RED) according to manufacturer’s instruction (Advanced Cell Diagnostics). The section slide was undergone H&E staining, followed by mounting it on a Tech-Med microscope slide using prolong gold mounting medium (Thomas Scientific).

### Transmission electron microscopy

Airway epithelium cultured on 6.5-mm Transwell membranes were fixed for 1 hour at RT in 2% PFA /2% glutaraldehyde in 0.1 M sodium cacodylate buffer (pH 7.4). The membranes were then removed from the inserts, rinsed with 0.1 M cacodylate buffer, postfixed on ice for 1 hour in 1% osmium tetroxide in 0.1 M cacodylate buffer containing 1.5% potassium ferrocyanide, dehydrated in a graded series of ethanol solutions (50%, 70%, 95%, and 100%), embedded in EMBed (Electron Microscopy Solutions, Inc.) and sectioned at 80-nm thickness in a direction parallel to the Transwell membrane. Thin sections were stained with 2% alcoholic uranyl acetate and 0.2% lead citrate and imaged using a Hitachi H-7500 transmission electron microscope.

### Cell area and cell perimeter measurements

The cell area and perimeter were determined for both RSV-GFP- and mock-infected cells via Fiji [45] with the plugin PaCeQuant [46] with a slight modification. Briefly, confocal images of the mock-infected cells were input into the Fiji program and converted into 8-bit grayscale images. The images were then input into the PaCeQuant plugin from the MiToBo plugin set, which segmented the images into cells and calculated the cell area and perimeter in multiple random areas, covering most cells of the image (at least 36 random cells). The RSV-GFP-infected cells were subjected to the same process as the mock-infected cells, with some differences. Whereas the mock-infected cell data were generated directly from the PaCeQuant plugin, the RSV-GFP-infected cells were analyzed only using the segmentation feature of PaCeQuant for successful demarcation of infected cells. The data for the RSV-infected cells, which were defined as those exhibiting GFP fluorescence, were then extracted from the segmented images of GFP-positive cells via the Magic Wand tool and the Region of Interest. The data represent at least two independent experiments.

### Epithelial layer thickness measurement

The airway epithelial layer thickness was evaluated using Imaris software. Confocal images (Z-stack, 60x oil objective) of the mock-infected or RSV-GFP-infected epithelium was evaluated for thickness using the 3D View tab in the IMARIS software program. The visible apical surface was stained for F-actin with rhodamine phalloidin (Texas Red), and the visible basal surface (bottom) was stained with DAPI. Height was then measured from the merged image (F-actin and DAPI) using the Magic Wand option. Briefly, an Ortho Slicer tab was created and set to the XZ plane. After rotating the slice into view and selecting a desired slice in the Ortho Slicer tab, a Measurement Points tab was created. In the Edit tab, the “intersects with” option was changed to “surface of object”, and in the setting tab, the line mode was changed to “pairs.” After switching back to the Edit tab of Measurement Points, two points were placed by holding the shift button on the keyboard and selecting a point on the apical layer and a point on the basal layer that were presumed to be in line on the X plane. To ensure that the two selected points were directly in line on the X plane, the XYZ position that appeared when one of the points was selected was altered, if necessary, to match the other point’s X position. The line between the two points automatically calculated a distance in microns. This generated 5 Z thickness values for each image. At least two images were quantified per treatment from each experiment. The data were then plotted in GraphPad Prism to compare epithelial height between mock-infected and RSV-infected epithelium. To determine the thickness of only GFP-infected cells, the above method was repeated for only the RSV-GFP-infected images, but rather than choosing random slices and apical and basal points, the measurements were intentionally chosen to include only infected epithelial areas. To do this, the FITC channel, which represented the RSV-GFP-infected cells, was turned on along with the DAPI and Texas Red channel. The slices and apical and basal points were selected to directly measure the epithelial height of the tissues at the locations of the RSV-GFP-infected cells.

### F-actin/G-actin ratio determination

Protein samples from ALI cultures were collected at 6 DPI and infected with RSV-GFP at an MOI of 4. Briefly, the apical part of a Transwell was washed twice with 1xPBS, and then all the cells were scraped out and transferred into a 1.5-ml tube. The cells were pelleted by centrifugation at 10,000 rpm for 5 min at 4°C. After removal of the supernatant, 110 μl of LAS2 buffer was prepared according to the protocol of a G-actin/F-actin In Vivo Assay Biochem Kit (Cytoskeleton, Inc.) and mixed with the pellet. The mixture was transferred into a QIAshredder tube (Qiagen) for centrifugation at 15,000 rpm for 3 min at 4°C in a tabletop centrifuge and then incubated at 37°C for 10 min. Then, 100 μl of the eluate was transferred into a Beckman 11-mm ultracentrifuge tube (Beckman Coulter) for ultracentrifugation in an Optima TLX ultracentrifuge (Beckman Coulter) with a TLA120.2 rotor (Beckman Coulter) at 60,000 rpm for 1 hour (37°C). 100 μl of supernatant was transferred into a new tube, and 100 μl of F-actin depolymerization buffer was added and mixed with the pellet. Both tubes were kept on ice for 1 hour, and then 25 μl of 5x SDS was added to each tube. Equal amounts of supernatant and pellet samples (15 μl) and G-actin protein standards (10, 50, and 100 μg) were separated on 4-12% Bis-tris SDS polyacrylamide gels, and the separated proteins were transferred by dry blotting onto polyvinylidene fluoride (PVDF) membranes according to the manufacturer’s instructions (Thermo Fisher Scientific). After transfer, the membranes were blocked in TBST/5% nonfat milk (10 mM Tris HCl pH 8.0, 150 mM NaCl, and 0.01% Tween 20/5% nonfat milk) for 30 min at RT and then washed in TBST 3 times for 10 min each per wash at RT. After blocking and washing, the membranes were incubated with the primary Ab (anti-actin rabbit polyclonal Ab) supplied with the G-actin/F-actin In Vivo Assay Biochem Kit (diluted 1:500 in TBST/0.1% nonfat milk) for 1 hour at RT and then washed 3 times in TBST for 10 min per wash at RT. The membranes were incubated with a secondary goat anti-rabbit HRP Ab (Sigma-Aldrich) diluted 1:5,000 in TBST/0.1% nonfat milk for 1 hour at RT and then washed 5 times in TBST for 10 min per wash at RT. The proteins were detected with a chemiluminescent substrate according to the manufacturer’s instructions (GE Healthcare). The membranes were processed for chemiluminescence detection of actin (43 kDa) using an Azure C600 system (Azure Biosystems, USA).

### Live-cell imaging

Airway epithelium cultured on 6.5-mm Transwell membranes was infected with RSV-GFP (MOI=4) at the apical side (described above). At 5 DPI, live-cell SiR-actin (Cytoskeleton, Inc.) was added to the complete medium at the basal side, and the epithelium was cultured for six hours before imaging using a 20x objective under a Leica DMi8 inverted fluorescence microscope equipped with an environmentally controlled chamber to maintain 37°C and 5% CO_2_. Images were taken every 5 min to capture the epithelial tissue (phase-contrast), RSV-GFP (AF488), and F-actin (Cy5) for approximately 16 hours. LASX software (Leica Microsystems) installed in the microscope was used to create a movie file.

### Membrane permeability assay

The differentiated epithelial layer permeability was determined by measuring TEER. TEER was measured using an epithelial volt-ohm meter (EVOM2, World Precision Instruments, Inc.). The EVOM2 was calibrated according to the manufacturer’s instructions, and the STX2 electrode was sterilized with 70% ethanol before use. The internal electrode (smaller in size) was placed in the apical part of each Transwell (PBS was added during TEER reading), and the external electrode (larger in size) was placed in the basal part of the Transwell, which contained ALI basal medium, to measure the membrane voltage and resistance of the epithelial layer. An empty Transwell insert (filled with 1xPBS) containing no NHBE cells was used to correct for background resistance. Three readings were taken for each Transwell. The TEER value of each sample was calculated by subtracting the background value.

### Ciliary function assay

Ciliary beating on the apical surfaces of cells in the differentiated epithelial layer was quantified by determining the CBF. Briefly, cilia were visualized in phase-contrast mode with a 20x objective on a Leica DMi8 microscope. For each Transwell, 6 different random fields were recorded for 2.1 seconds at 120 frames per second. The images were captured at 37°C and analyzed using the SAVA system V.2.1.15 to determine the CBF in Hz.

### Western blotting

Protein samples were collected from the airway epithelium infected with RSV-GFP at an MOI of 4 at 6 DPI. Briefly, the apical part of a Transwell was washed 2 times with 1xPBS and then scraped to collect all the cells, which were transferred into a 1.5-ml tube. The cells were pelleted by centrifugation at 10,000 rpm for 5 min at 4°C. After removal of the supernatant, the cells were collected into a QIAshredder tube and incubated for 1 min at RT with 100 μl of gel-loading buffer made from 2.5 ml of 4x LDS loading buffer (Thermo Fisher Scientific), 1 proteinase inhibitor tablet and 7.5 ml of PBS. The cell mixture was centrifuged at 15,000 rpm for 3 min at 4°C. The eluate from the QIAshredder tube was stored at −20°C in a freezer. A similar approach was used to collect protein samples from NHBE monolayers in a 24-well plate. The protein concentrations were measured using a BCA protein assay kit (Thermo Fisher Scientific). To detect ARP2, total protein (10 μg) was separated on 4-12% Bis-tris SDS polyacrylamide gels (nonreducing), and the separated proteins were transferred by dry blotting onto PVDF membranes according to the manufacturer’s instructions (Life Technologies). Similarly, to detect NS2, we run 10 μl of apical wash of RSV-GFP-infected transwell on a 4-12% Bis-tris SDS polyacrylamide gels (nonreducing and nondenaturing). For E-cadherin, Tricellulin (MARVELD2), MUC5B, respiratory syncytial virus nucleoprotein, respiratory syncytial virus phosphoprotein detection, samples were denatured at 90°C for 10 min with a 10x reducing agent (Thermo Fisher Scientific) before electrophoresis on 4-12% Bis-tris gels. α-tubulin and GAPDH were used as loading control. For all analyses, the PVDF membranes were incubated in LI-COR blocking buffer (1:1 in 1x PBS) (LI-COR Biosciences) for 1 hour at RT and then overnight at 4°C with a primary Ab (1:1,000) in LI-COR blocking buffer. The membranes were washed 3 times for 5 min with PBS and then incubated with a secondary Ab (1:15,000) (LI-COR Biosciences) for 1 hour at RT. After washing 3 times for 5 min with 1xPBS, fluorescence was analyzed using an Odyssey imaging system (LI-COR Biosciences). To detect ARP2, E-cadherin, and MUC5B, rabbit monoclonal antibodies including anti-ARP2 (Cat# ab129018, Abcam), anti-E-cadherin (Cat# 3195S, Cell Signaling Technologies), anti-MUC5B (1:500) (Cat# HPA008246, Atlas Antibodies), anti-NS2 (rabbit mAb, custom made by ABclonal) were used, respectively followed by a goat anti-rabbit IRDye800-labeled secondary Ab (Cat# 926-32211, LI-COR Biosciences). To detect GAPDH and Tricellulin (MARVELD2), rabbit polyclonal antibodies including anti-GAPDH (Cat# G9545, Sigma-Aldrich) and anti- MARVELD2 (Cat# 48-8400, Thermo Fisher Scientific), were used, respectively followed by a goat anti-rabbit IRDye800-labeled secondary Ab (Cat# 926-32211, LI-COR Biosciences). To detect α-tubulin, respiratory syncytial virus nucleoprotein and respiratory syncytial virus phosphoprotein, mouse monoclonal antibodies including anti- α tubulin (Cat# T6199, Sigma-Aldrich), anti-respiratory syncytial virus nucleoprotein (Cat# ab94806, Abcam) and anti-respiratory syncytial virus phosphoprotein (Cat# ab94965, Abcam) were used, respectively followed by a goat anti-mouse IRDye680-labeled secondary Ab (Cat# 926-68070, LI-COR Biosciences). Image Studio 5.2 (LI-COR Biosciences) was used to quantify protein signal.

### Cell cytotoxicity determination

40 μl of apical wash (1x PBS) and basal medium from airway epithelia mock-infected or infected with RSV-GFP (MOI=4) at 6 DPI were collected and assessed for cell cytotoxicity with a commercially available Cell Cytotoxicity Detection Kit (LDH). 40 μl of sample was mixed with 40 μl of freshly prepared reaction mixture and incubated for 30 min at RT. PBS and basal medium were used for the low control and for high control the cells were treated with 2% Triton X-100 (Sigma-Aldrich) for 30 min at room temperature, and the absorbance of the samples was measured at 490 nm by using a BioTek Synergy HT (BioTeK Instruments). The cytotoxicity percentage was calculated by using the following equation: cytotoxicity = (experimental value – low control/high control – low control) x 100.

### RNA extraction

Airway epithelium cultured on 6.5-mm Transwell membranes was washed and treated for 1 min at RT with RLT buffer (Qiagen) with 0.01 % β-mercaptoethanol (Sigma-Aldrich). The cells were scraped using a cell scraper, collected into a QIAshredder tube and centrifuged at 15,000 rpm for 3 min at 4°C. The eluate was used to extract total RNA using a Total DNA/RNA Extraction Kit (Qiagen), and DNase I treatment was performed to remove DNA from the samples according to the manufacturer’s instructions. The RNA concentration was determined with an Epoch instrument (BioTek Instruments).

### RNA-seq for transcriptome analysis

RNA was extracted (as described above) from ALI cultures at 6 DPI after mock infection or infection with RSV-WT at an MOI of 4. Preliminary quality control analysis of the FASTQ files was performed with FastQC v0.11.8 [98]. The adapters were trimmed using Trimmomatic v0.39 [99]. The reads were aligned to the human reference genome (hg19) with STAR v2.7.1a [59]. Gene expression was quantified using CuffNorm v2.2.1 [100]. The read counts were summarized using featureCounts v1.4.6 [101]. Differential expression analysis of RSV- vs mock-infected samples was performed using the R/Bioconductor package DESeq2 v1.24.0 [102]. A gene was considered differentially expressed if it had an FDR of 0.05 or less. GO and KEGG pathway enrichment analyses were performed using the R/Bioconductor package clusterProfiler v3.12.0 [103]. A GO term or KEGG pathway was considered enriched if it had a p-value of 0.05 or less, a Benjamini-Hochberg adjusted p-value of 0.05 or less and a q-value of 0.05 or less.

### siRNA transfection and RSV infection

NHBE cells were transfected with siARP2 (s223082, Thermo Fisher Scientific) or a nonspecific siControl (Silencer Select Negative Control #2, Thermo Fisher Scientific). Briefly, 2×10^4^ cells were seeded in PureCol-coated 24-well plates and incubated overnight at 37°C in a CO_2_ incubator. The cells reached approximately 70% confluency the next day. The cells were transfected with an siRNA transfection mixture containing 100 μl of complete AEC medium, 1.5 μl of RNAiMax transfection reagent (Thermo Fisher Scientific), and 7.5 μl of siRNA (2 μM) for 48 hours at 37°C. Over 80% ARP2 knockdown was achieved, as confirmed by Western blotting. For RSV-WT infection, the medium was removed, and 100 μl of virus inoculum (prepared in complete AEC medium) (RSV-WT, MOI=0.5) was added to the NHBE monolayer. The mixture was incubated for one hour at 37°C. The virus inoculum was then removed from the well, and the cells were washed twice with complete AEC medium and incubated in complete AEC medium at 37°C in a CO_2_ incubator for 72 hours.

### Cytokine and chemokine measurements

We collected samples from the apical and basal parts of mock-infected and RSV-GFP-infected (MOI = 4) respiratory epithelial cultures at 2 hours, 6 hours, 12 hours and 24 hours post infection. Apical washes and basal medium were collected to determine chemokine induction in the epithelium. To analyze the concentrations of cytokines (such as TNF-α) and chemokines (such as IP10/CXCL10), a LegendPlex Human Anti-Virus Response Panel were used according to the manufacturer’s directions (BioLegend). Briefly, 20 μl of supernatant from each sample was added individually to a well of a 96-well V-bottom plate. 10 μl of analyte beads were added to each well, and the plate was incubated at RT with shaking at 800 rpm on a plate shaker for two hours while covered in aluminum foil. After incubation, the beads were washed twice for 5 min per wash by centrifugation at ~250x *g* at RT with the included wash buffer diluted to 1x in Milli-Q deionized water. After the second wash, the beads were resuspended by shaking at 800 rpm for 1 min, and then biotinylated Abs were added to each well. The plate was shaken at 800 rpm for one hour at RT while covered in aluminum foil. After one hour, 10 μl of SA-PE provided with the kit (BioLegend) was added to each well, and the plate was again shaken at 800 rpm while covered in aluminum foil for 30 min at RT. The beads were then washed twice with 1x wash buffer and run on a BD FACS Symphony flow cytometer (BD Biosciences) under the default settings by following the manufacturer’s instructions (BioLegend). The data were analyzed using LegendPlex software (BioLegend), and the concentration of IP10/CXCL10 was determined by comparing the mean fluorescence intensity of each sample supernatant to its respective standard curve, which was generated by following the manufacturer’s instructions (BioLegend). Data were analyzed using the LegendPlex data analysis software v.8 for Windows.

### Cell viability assay

NHBE cell monolayer viability was evaluated using a resazurin-based alamarBlue assay (Thermo Fisher Scientific) according to the manufacturer’s protocol. Briefly, a 10% volume of alamarBlue was added to the cell culture medium and incubated at 37°C for 3-4 hours. To evaluate cell viability, alamarBlue fluorescence, a marker for metabolic activity, was analyzed using a Synergy 2 Multi Mode microplate reader (BioTek).

### RSV transcript quantification

To quantify transcript expression in RSV-infected cells, reads were aligned to the human RSV genome (GenBank accession KT992094) with STAR v2.7.1a [59]. The read counts were summarized using featureCounts v1.4.6 [101]. The FPKM values were calculated using the fpkm function in DESeq2 v1.24.0 [102].

### Statistical analysis

Parameters such as the number of replicates, number of independent experiments, mean+/− Standard error of mean (SEM), and statistical significance are reported in the figures and figure legends. A p-value less than 0.05 was considered to indicate significance. Where appropriate, the statistical tests and post hoc statistical analysis methods are noted in the figure legends or Methods section.

## Supporting Information

**Fig S1. Bronchial airway epithelium from primary NHBE cells using 3D cell culture (air-liquid interface culture).** (A) A schematic illustration of the establishment of an airway epithelium model with primary NHBE cells. (B) The whole-genome RNA-seq data from P1, P2, P3, and P4 cells were compared for gene expression analysis. The correlations are shown as a heatmap with the correlation values plotted. (C) Four-week-differentiated P4 NHBE cells (airway epithelium) were fixed, permeabilized, and stained with an anti-acetyl-α-tubulin primary Ab (specific for cilium), an anti-MUC5AC primary Ab (specific for mucin) or an anti-MUC5B primary Ab (specific for mucin) and respective secondary Abs conjugated with either AF647 (shown in cyan) or AF488 (shown in green). F-actin was detected by rhodamine phalloidin staining. Ciliated epithelial cells and goblet cells were identified by confocal microscopy using a 60x objective. F-actin is shown in red, and other proteins are shown in cyan (MUC5AC) or green (MUC5B) in the images. The scale bar is 5 μm. (D) Adherens junctions, tight junctions, and tricellular junctions were identified by using anti-E-cadherin, anti-ZO-1, and anti-tricellulin primary Abs, and the respective secondary Abs conjugated with AF647 (shown in cyan) were identified by confocal microscopy using a 60**X** objective. F-actin is shown in red, and the other proteins are shown in cyan in the images. The scale bar is 5 μm.

**Fig S2: RSV infection in Airway epithelium.** (A). The airway epithelium was mock-infected or infected with RSV-WT at an MOI of 4. At six days post infection (DPI), the RSV-infected airway epithelium was fixed and sectioned for transmission electron microcopy imaging. A representative image of the RSV-WT infected airway epithelium at magnification (30,000x). “*”, “#”, and “^” indicate disorganization of granules in goblet cells, the appearance of interdigitating elements, and an increased density of microvilli, respectively. Two independent filamentous RSV virions that were budded out from the infected-ciliated cells are showing in the right panel and their magnified images are shown in the inset. (B). Formalin fixed paraffin-embedded (FFPE) RSV-infected airway epithelium (6 DPI, MOI=4) was stained for RSV N mRNA by using specific probe targeting N mRNA using RNAscope. Scale bar 25 nm.

**Fig S3. RSV infects ciliated cells and expands the actin cytoskeleton**. (A) At 3 DPI, RSV-WT-infected (MOI = 0.5) airway epithelium was fixed, permeabilized, and stained (via immunofluorescence) for the RSV F protein (anti-F antibody, green) and cilia (acetyl-α- tubulin, cyan). F-actin (red) and nuclei (blue) were detected by staining with rhodamine phalloidin and DAPI, respectively. The scale bar is 5 μm. The arrowhead indicates a filamentous RSV virion. (B and C) Four-week-differentiated NHBE cells (airway epithelium) were mock-infected or infected with either RSV-WT (A2 strain), HMPV, or influenza A virus (PR8 strain) at an MOI of 0.5 for 3 days. The epithelium was fixed, permeabilized, and stained with an anti-acetyl-α--tubulin Ab (specific for cilia) (shown in cyan) and respective viral protein-specific Abs (anti-RSV F, anti-HMPV N, and anti-influenza NP Abs) (shown in green). F-actin (shown in red) and nuclei (shown in blue) were detected by staining with rhodamine phalloidin and DAPI, respectively. (D) SARS-CoV-2 (top image) and MERS (bottom image) were infected airway epithelium at MOI = 0.1 for four days [35]. The epithelium was fixed, permeabilized, and stained with an anti-acetyl-α--tubulin Ab (specific for cilia) (shown in cyan) and respective viral protein-specific Abs (anti-SARS-CoV-2 N and anti-MERS N Abs) (shown in green). F-actin (shown in red) and nuclei (shown in blue) were detected by staining with rhodamine phalloidin and DAPI, respectively. The images were taken by a confocal microscope using a 60**x** oil objective. The scale bar is 15 μm for B, 20 μm for C, and 15 μm for D.

**Fig S4. Expanded actin cytoskeleton in RSV-infected cells**. (A) 3D image of an RSV-GFP-infected (MOI = 4) airway epithelium at 6 DPI. F-actin (detected by phalloidin staining) is shown in red, and RSV-GFP-infected cells are shown in green. The 3D image was generated in Imaris 9 (Bitplane) using Z-stack confocal images. The scale bar is 20 μm. (B) Representative images showing the process followed for confocal image quantification to determine the cell area and perimeter (FIG 1). Mock-infected or RSV-GFP-infected airway epithelium was processed for mosaic transformation into 8-bit grayscale images with Fiji [45], after which the PaCeQuant plugin [46] was used in Fiji.

**Fig S5. RSV infection does not change adherin-junctions, tight-junctions, or tricellular junctions.** (A) Mock- and RSV-GFP-infected (MOI = 4) airway epithelium was stained with primary Abs against E-cadherin, ZO-1 and tricellulin and respective secondary Abs conjugated with AF647 (shown in cyan) at 6 DPI. F-actin (shown in red) and nuclei (shown in blue) were stained with rhodamine phalloidin and DAPI, respectively. The scale bar is 20 μm. (B and C) Western blot-based detection of E-cadherin and tricellulin in mock- and RSV-GFP-infected (MOI = 4) epithelium at 6 DPI. Here, α-tubulin was also detected as a loading control.

**Fig S6. RSV induces transcriptional changes.** (A)PCA of the transcriptome data. The data for three individual samples of mock- and RSV-WT-infected (MOI = 4, at 6 DPI) epithelium are plotted. (B) Transcriptome data from mock- and RSV-WT-infected (MOI = 4, at 6 DPI) epithelium overlaid with the “regulation of the actin cytoskeleton” KEGG pathway. The genes are colored according to their degree of expression. Green represents downregulated genes, red represents upregulated genes, and white represents unchanged genes. (C) The top-ranked differentially expressed genes are overlaid with the “regulation of actin-based motility by Rho pathway” from Ingenuity Pathway Analysis (IPA) (Qiagen). The genes are colored according to their p-values.

**Fig S7. RSV modulates TNF-α signaling.** Transcriptome data from mock- and RSV-GFP-infected (MOI = 4, at 6 DPI) epithelium were overlaid with the “TNF-α signaling” KEGG pathway. The genes are colored according to their degrees of expression. Green represents downregulated genes, red represents upregulated genes, and white represents unchanged genes.

**Fig S8. Robust N and P production in the infected respiratory epithelium.** (A) RSV N levels in RSV-GFP-infected (MOI = 4) epithelium were detected by Western blotting. For the loading control, GAPDH was detected. The membrane was reprobed to detect RSV P. (B) RSV N and P levels were measured by normalization with the GAPDH level.

**Table S1. List of significantly modulated (differentially expressed) genes.**

**Table S2. List of the top 86 GO biological processes associated with the differentially expressed genes.**

**Table S3. List of the differentially expressed genes associated with actin regulation.**

**Table S4. List of the top 63 KEGG pathway biological processes associated with the differentially expressed genes.**

**Table S5. List of the differentially expressed genes in the TNF signaling pathway.**

**Movie S1. RSV-GFP infection of the airway epithelium.** An RSV-GFP-infected (MOI=4) respiratory epithelium was imaged at 5 DPI for around 15 hours. The cells were imaged every 5 min. Time is presented in hours:min:sec. Green represents RSV-GFP-infected cells. The movie file is representative at least five independent positions.

**Movie S2. RSV-GFP infection of the airway epithelium.** An RSV-GFP-infected (MOI=4) respiratory epithelium was imaged at 5 DPI for around 15 hours. The cells were imaged every 5 min. Time is presented in hours:min:sec. Green and purple represent RSV-GFP-infected cells and F-actin (SiR-Actin, live cell dye), respectively.

## Acknowledgments

This work was funded by NIH/NIGMS P20GM113123. We thank Dr. Nadeem Khan, UND, for providing the influenza A virus PR8 strain. We thank the Microscopy Core (UND, Grand Forks) funded by NIH P20GM103442 of the INBRE program for providing access to an Olympus FV300 confocal microscope and Dr Swojani Shrestha for technical support. We also thank the Imaging Core (UND, Grand Forks) funded by NIH P20GM113123, NIH U54GM128729, and UNDSMHS for providing access to the TEM imaging system and Imaris image analysis software. We acknowledge the Genomics core (UND, Grand Forks) for RNA-seq funded by U54GM128729 and 2P20GM104360-06A1.

## Genomic data submission

The RNA-seq data are available through the GEO under the accession number GSE146795.

## Author contributions

Conceptualization: M.M., S.N.T.

Format Analysis: M.M., J.K.O., M.K., B.D.K., D.P., S.Y.

Investigations: M.M., S.N.T., J.K.O., L.H.

Resources: K.L.B.

Methodology: M.M., S.N.T, B.D.K., D.P.

Supervisions: M.M.

Visualization: S.N.T., K.R., S.D., B.G.

Writing - original draft: M.M., S.N.T., J.K.O.

Writing - review and editing: M.M., S.N.T., J.K.O.

## References

1. Ackerson B, Tseng HF, Sy LS, Solano Z, Slezak J, Luo Y, et al. Severe Morbidity and Mortality Associated With Respiratory Syncytial Virus Versus Influenza Infection in Hospitalized Older Adults. Clin Infect Dis. 2019;69(2):197–203. Epub 2018/11/20. doi: 10.1093/cid/ciy991. PubMed PMID: 30452608; PubMed Central PMCID: PMCPMC6603263.

2. Hall CB, Simoes EA, Anderson LJ. Clinical and epidemiologic features of respiratory syncytial virus. Curr Top Microbiol Immunol. 2013;372:39–57. Epub 2013/12/24. doi: 10.1007/978-3-642-38919-1_2. PubMed PMID: 24362683.

3. Sommer C, Resch B, Simoes EA. Risk factors for severe respiratory syncytial virus lower respiratory tract infection. Open Microbiol J. 2011;5:144–54. Epub 2012/01/21. doi: 10.2174/1874285801105010144. PubMed PMID: 22262987; PubMed Central PMCID: PMCPMC3258650.

4. Falsey AR, Walsh EE. Respiratory syncytial virus infection in adults. Clin Microbiol Rev. 2000;13(3):371–84. Epub 2000/07/25. doi: 10.1128/cmr.13.3.371-384.2000. PubMed PMID: 10885982; PubMed Central PMCID: PMCPMC88938.

5. Welliver TP, Garofalo RP, Hosakote Y, Hintz KH, Avendano L, Sanchez K, et al. Severe human lower respiratory tract illness caused by respiratory syncytial virus and influenza virus is characterized by the absence of pulmonary cytotoxic lymphocyte responses. J Infect Dis. 2007;195(8):1126–36. Epub 2007/03/16. doi: 10.1086/512615. PubMed PMID: 17357048; PubMed Central PMCID: PMCPMC7109876.

6. Aherne W, Bird T, Court SD, Gardner PS, McQuillin J. Pathological changes in virus infections of the lower respiratory tract in children. J Clin Pathol. 1970;23(1):7–18. Epub 1970/02/01. doi: 10.1136/jcp.23.1.7. PubMed PMID: 4909103; PubMed Central PMCID: PMCPMC474401.

7. Carvajal JJ, Avellaneda AM, Salazar-Ardiles C, Maya JE, Kalergis AM, Lay MK. Host Components Contributing to Respiratory Syncytial Virus Pathogenesis. Front Immunol. 2019;10:2152. Epub 2019/10/02. doi: 10.3389/fimmu.2019.02152. PubMed PMID: 31572372; PubMed Central PMCID: PMCPMC6753334.

8. Kim MC, Kim MY, Lee HJ, Lee SO, Choi SH, Kim YS, et al. CT findings in viral lower respiratory tract infections caused by parainfluenza virus, influenza virus and respiratory syncytial virus. Medicine (Baltimore). 2016;95(26):e4003. Epub 2016/07/02. doi: 10.1097/MD.0000000000004003. PubMed PMID: 27368011; PubMed Central PMCID: PMCPMC4937925.

9. Osborne D. Radiologic appearance of viral disease of the lower respiratory tract in infants and children. AJR Am J Roentgenol. 1978;130(1):29–33. Epub 1978/01/01. doi: 10.2214/ajr.130.1.29. PubMed PMID: 202157.

10. Darras KE, Roston AT, Yewchuk LK. Imaging Acute Airway Obstruction in Infants and Children. Radiographics. 2015;35(7):2064–79. Epub 2015/10/27. doi: 10.1148/rg.2015150096. PubMed PMID: 26495798.

11. Colby TV. Bronchiolitis: pathologic considerations. American journal of clinical pathology. 1998;109(1):101–9.

12. Ryu JH, Myers JL, Swensen SJ. Bronchiolar disorders. Am J Respir Crit Care Med. 2003;168(11):1277–92. Epub 2003/12/04. doi: 10.1164/rccm.200301-053SO. PubMed PMID: 14644923.

13. Colby TV. Bronchiolitis. Pathologic considerations. Am J Clin Pathol. 1998;109(1):101–9. Epub 1998/01/14. doi: 10.1093/ajcp/109.1.101. PubMed PMID: 9426525.

14. Hardy KA, Schidlow DV, Zaeri N. Obliterative bronchiolitis in children. Chest. 1988;93(3):460–6.

15. Visscher DW, Myers JL. Bronchiolitis: the pathologist’s perspective. Proceedings of the American Thoracic Society. 2006;3(1):41–7.

16. Hall CB. Respiratory syncytial virus and parainfluenza virus. New England journal of medicine. 2001;344(25):1917–28.

17. Grydeland TB, Dirksen A, Coxson HO, Eagan TM, Thorsen E, Pillai SG, et al. Quantitative computed tomography measures of emphysema and airway wall thickness are related to respiratory symptoms. American journal of respiratory and critical care medicine. 2010;181(4):353–9.

18. Openshaw PJ, Dean GS, Culley FJ. Links between respiratory syncytial virus bronchiolitis and childhood asthma: clinical and research approaches. The Pediatric infectious disease journal. 2003;22(2):S58–S65.

19. Wilkinson TM, Donaldson GC, Johnston SL, Openshaw PJ, Wedzicha JA. Respiratory syncytial virus, airway inflammation, and FEV1 decline in patients with chronic obstructive pulmonary disease. American journal of respiratory and critical care medicine. 2006;173(8):871–6.

20. Sikkel MB, Quint JK, Mallia P, Wedzicha JA, Johnston SL. Respiratory syncytial virus persistence in chronic obstructive pulmonary disease. The Pediatric infectious disease journal. 2008;27(10):S63–S70.

21. Hendricks DA, Baradaran K, McIntosh K, Patterson JL. Appearance of a soluble form of the G protein of respiratory syncytial virus in fluids of infected cells. Journal of General Virology. 1987;68(6):1705–14.

22. Walsh E, Hruska J. Monoclonal antibodies to respiratory syncytial virus proteins: identification of the fusion protein. Journal of Virology. 1983;47(1):171–7.

23. Talukdar SN MM. Respiratory Syncytia Virus 2022. In: RNA Viruses Infection [Internet]. Intech Open.

24. Spann KM, Tran K-C, Chi B, Rabin RL, Collins PL. Suppression of the induction of alpha, beta, and gamma interferons by the NS1 and NS2 proteins of human respiratory syncytial virus in human epithelial cells and macrophages. Journal of virology. 2004;78(8):4363–9.

25. Spann KM, Tran KC, Collins PL. Effects of nonstructural proteins NS1 and NS2 of human respiratory syncytial virus on interferon regulatory factor 3, NF-κB, and proinflammatory cytokines. Journal of virology. 2005;79(9):5353–62.

26. Paluck A, Osan J, Hollingsworth L, Talukdar SN, Saegh AA, Mehedi M. Role of ARP2/3 Complex-Driven Actin Polymerization in RSV Infection. Pathogens. 2021;11(1):26.

27. Garcı́a J, Garcı́a-Barreno B, Vivo A, Melero JA. Cytoplasmic inclusions of respiratory syncytial virus-infected cells: formation of inclusion bodies in transfected cells that coexpress the nucleoprotein, the phosphoprotein, and the 22K protein. Virology. 1993;195(1):243–7.

28. Mehedi M, McCarty T, Martin SE, Le Nouen C, Buehler E, Chen YC, et al. Actin-Related Protein 2 (ARP2) and Virus-Induced Filopodia Facilitate Human Respiratory Syncytial Virus Spread. PLoS Pathog. 2016;12(12):e1006062. Epub 2016/12/08. doi: 10.1371/journal.ppat.1006062. PubMed PMID: 27926942; PubMed Central PMCID: PMCPMC5142808.

29. Mehedi M, Smelkinson M, Kabat J, Ganesan S, Collins PL, Buchholz UJ. Multicolor Stimulated Emission Depletion (STED) Microscopy to Generate High-resolution Images of Respiratory Syncytial Virus Particles and Infected Cells. Bio Protoc. 2017;7(17). doi: 10.21769/BioProtoc.2543. PubMed PMID: 29057295; PubMed Central PMCID: PMCPMC5650235.

30. Zhang L, Peeples ME, Boucher RC, Collins PL, Pickles RJ. Respiratory syncytial virus infection of human airway epithelial cells is polarized, specific to ciliated cells, and without obvious cytopathology. J Virol. 2002;76(11):5654–66. doi: 10.1128/jvi.76.11.5654-5666.2002. PubMed PMID: 11991994; PubMed Central PMCID: PMCPMC137037.

31. Villenave R, Thavagnanam S, Sarlang S, Parker J, Douglas I, Skibinski G, et al. In vitro modeling of respiratory syncytial virus infection of pediatric bronchial epithelium, the primary target of infection in vivo. Proc Natl Acad Sci U S A. 2012;109(13):5040–5. Epub 2012/03/14. doi: 10.1073/pnas.1110203109. PubMed PMID: 22411804; PubMed Central PMCID: PMCPMC3323997.

32. Rayner RE, Makena P, Prasad GL, Cormet-Boyaka E. Optimization of Normal Human Bronchial Epithelial (NHBE) Cell 3D Cultures for in vitro Lung Model Studies. Sci Rep. 2019;9(1):500. doi: 10.1038/s41598-018-36735-z. PubMed PMID: 30679531; PubMed Central PMCID: PMCPMC6346027.

33. Upadhyay S, Palmberg L. Air-Liquid Interface: Relevant In Vitro Models for Investigating Air Pollutant-Induced Pulmonary Toxicity. Toxicol Sci. 2018;164(1):21–30. Epub 2018/03/14. doi: 10.1093/toxsci/kfy053. PubMed PMID: 29534242.

34. Pawlina W. In: Pawlina W, editor. Histology a Text Book and Atlas with Correlated Cell and Molecualr Biology. 7th ed: Wolters Kluwer; 2016. p. 671–97.

35. Osan J, Talukdar SN, Feldmann F, DeMontigny BA, Jerome K, Bailey KL, et al. Goblet Cell Hyperplasia Increases SARS-CoV-2 Infection in Chronic Obstructive Pulmonary Disease. Microbiol Spectr. 2022:e0045922. Epub 20220713. doi: 10.1128/spectrum.00459-22. PubMed PMID: 35862971.

36. Holgate ST. Epithelium dysfunction in asthma. J Allergy Clin Immunol. 2007;120(6):1233–44; quiz 45-6. Epub 2007/12/13. doi: 10.1016/j.jaci.2007.10.025. PubMed PMID: 18073119.

37. Kojima T, Go M, Takano K, Kurose M, Ohkuni T, Koizumi J, et al. Regulation of tight junctions in upper airway epithelium. Biomed Res Int. 2013;2013:947072. Epub 2013/03/20. doi: 10.1155/2013/947072. PubMed PMID: 23509817; PubMed Central PMCID: PMCPMC3591135.

38. Liesman RM, Buchholz UJ, Luongo CL, Yang L, Proia AD, DeVincenzo JP, et al. RSV-encoded NS2 promotes epithelial cell shedding and distal airway obstruction. J Clin Invest. 2014;124(5):2219–33. Epub 2014/04/10. doi: 10.1172/JCI72948. PubMed PMID: 24713657; PubMed Central PMCID: PMCPMC4001550.

39. Yao X, Ireland SK, Pham T, Temple B, Chen R, Raj MH, et al. TLE1 promotes EMT in A549 lung cancer cells through suppression of E-cadherin. Biochemical and biophysical research communications. 2014;455(3-4):277–84.

40. Jumat MR, Yan Y, Ravi LI, Wong P, Huong TN, Li C, et al. Morphogenesis of respiratory syncytial virus in human primary nasal ciliated epithelial cells occurs at surface membrane microdomains that are distinct from cilia. Virology. 2015;484:395–411. Epub 2015/08/02. doi: 10.1016/j.virol.2015.05.014. PubMed PMID: 26231613.

41. de Graaf M, Herfst S, Aarbiou J, Burgers PC, Zaaraoui-Boutahar F, Bijl M, et al. Small hydrophobic protein of human metapneumovirus does not affect virus replication and host gene expression in vitro. PLoS One. 2013;8(3):e58572. Epub 2013/03/14. doi: 10.1371/journal.pone.0058572. PubMed PMID: 23484037; PubMed Central PMCID: PMCPMC3590193.

42. Matrosovich MN, Matrosovich TY, Gray T, Roberts NA, Klenk HD. Human and avian influenza viruses target different cell types in cultures of human airway epithelium. Proc Natl Acad Sci U S A. 2004;101(13):4620–4. Epub 2004/04/09. doi: 10.1073/pnas.0308001101. PubMed PMID: 15070767; PubMed Central PMCID: PMCPMC384796.

43. Lee RJ, Kofonow JM, Rosen PL, Siebert AP, Chen B, Doghramji L, et al. Bitter and sweet taste receptors regulate human upper respiratory innate immunity. J Clin Invest. 2014;124(3):1393–405. Epub 20140217. doi: 10.1172/JCI72094. PubMed PMID: 24531552; PubMed Central PMCID: PMCPMC3934184.

44. Lee RJ, Xiong G, Kofonow JM, Chen B, Lysenko A, Jiang P, et al. T2R38 taste receptor polymorphisms underlie susceptibility to upper respiratory infection. J Clin Invest. 2012;122(11):4145–59. Epub 20121008. doi: 10.1172/JCI64240. PubMed PMID: 23041624; PubMed Central PMCID: PMCPMC3484455.

45. Schindelin J, Arganda-Carreras I, Frise E, Kaynig V, Longair M, Pietzsch T, et al. Fiji: an open-source platform for biological-image analysis. Nat Methods. 2012;9(7):676–82. Epub 2012/06/30. doi: 10.1038/nmeth.2019. PubMed PMID: 22743772; PubMed Central PMCID: PMCPMC3855844.

46. Moller B, Poeschl Y, Plotner R, Burstenbinder K. PaCeQuant: A Tool for High-Throughput Quantification of Pavement Cell Shape Characteristics. Plant Physiol. 2017;175(3):998–1017. Epub 2017/09/22. doi: 10.1104/pp.17.00961. PubMed PMID: 28931626; PubMed Central PMCID: PMCPMC5664455.

47. Hayden MS, Ghosh S. Regulation of NF-kappaB by TNF family cytokines. Semin Immunol. 2014;26(3):253–66. Epub 2014/06/25. doi: 10.1016/j.smim.2014.05.004. PubMed PMID: 24958609; PubMed Central PMCID: PMCPMC4156877.

48. Khalfan M. Gene Set Enrichment Analysis with ClusterProfiler 2020 [cited 2020 09/22/2020]. Available from: https://learn.gencore.bio.nyu.edu/rna-seq-analysis/gene-set-enrichment-analysis/.

49. Mehedi M, Collins PL, Buchholz UJ. A novel host factor for human respiratory syncytial virus. Communicative & integrative biology. 2017;10(3):e1006062.

50. Paluck A, Osan J, Hollingsworth L, Talukdar SN, Saegh AA, Mehedi M. Role of ARP2/3 Complex-Driven Actin Polymerization in RSV Infection. Pathogens. 2021;11(1). Epub 20211226. doi: 10.3390/pathogens11010026. PubMed PMID: 35055974; PubMed Central PMCID: PMCPMC8781601.

51. Goley ED, Welch MD. The ARP2/3 complex: an actin nucleator comes of age. Nat Rev Mol Cell Biol. 2006;7(10):713–26. Epub 2006/09/23. doi: 10.1038/nrm2026. PubMed PMID: 16990851.

52. Garcia-Gonzalez J, Kebrlova S, Semerak M, Lacek J, Kotannal Baby I, Petrasek J, et al. Arp2/3 Complex Is Required for Auxin-Driven Cell Expansion Through Regulation of Auxin Transporter Homeostasis. Front Plant Sci. 2020;11:486. Epub 2020/05/20. doi: 10.3389/fpls.2020.00486. PubMed PMID: 32425966; PubMed Central PMCID: PMCPMC7212389.

53. Gournier H, Goley ED, Niederstrasser H, Trinh T, Welch MD. Reconstitution of human Arp2/3 complex reveals critical roles of individual subunits in complex structure and activity. Mol Cell. 2001;8(5):1041–52. Epub 2001/12/14. doi: 10.1016/s1097-2765(01)00393-8. PubMed PMID: 11741539.

54. Ivanov AI. Actin motors that drive formation and disassembly of epithelial apical junctions. Front Biosci. 2008;13:6662–81. Epub 2008/05/30. doi: 10.2741/3180. PubMed PMID: 18508686.

55. Laing AG, Lorenc A, Del Molino Del Barrio I, Das A, Fish M, Monin L, et al. A dynamic COVID-19 immune signature includes associations with poor prognosis. Nat Med. 2020;26(10):1623–35. Epub 2020/08/19. doi: 10.1038/s41591-020-1038-6. PubMed PMID: 32807934.

56. Hamana A, Takahashi Y, Uchida T, Nishikawa M, Imamura M, Chayama K, et al. Evaluation of antiviral effect of type I, II, and III interferons on direct-acting antiviral-resistant hepatitis C virus. Antiviral Res. 2017;146:130–8. Epub 2017/09/03. doi: 10.1016/j.antiviral.2017.08.017. PubMed PMID: 28864074.

57. Liu T, Zhang L, Joo D, Sun SC. NF-kappaB signaling in inflammation. Signal Transduct Target Ther. 2017;2. Epub 2017/11/22. doi: 10.1038/sigtrans.2017.23. PubMed PMID: 29158945; PubMed Central PMCID: PMCPMC5661633.

58. Osborn L, Kunkel S, Nabel GJ. Tumor necrosis factor alpha and interleukin 1 stimulate the human immunodeficiency virus enhancer by activation of the nuclear factor kappa B. Proc Natl Acad Sci U S A. 1989;86(7):2336–40. Epub 1989/04/01. doi: 10.1073/pnas.86.7.2336. PubMed PMID: 2494664; PubMed Central PMCID: PMCPMC286907.

59. Dobin A, Davis CA, Schlesinger F, Drenkow J, Zaleski C, Jha S, et al. STAR: ultrafast universal RNA-seq aligner. Bioinformatics. 2013;29(1):15–21. Epub 20121025. doi: 10.1093/bioinformatics/bts635. PubMed PMID: 23104886; PubMed Central PMCID: PMCPMC3530905.

60. Rayner RE, Wellmerling J, Osman W, Honesty S, Alfaro M, Peeples ME, et al. In vitro 3D culture lung model from expanded primary cystic fibrosis human airway cells. J Cyst Fibros. 2020;19(5):752–61. Epub 2020/06/23. doi: 10.1016/j.jcf.2020.05.007. PubMed PMID: 32565193.

61. Osan JK, DeMontigny BA, Mehedi M. Immunohistochemistry for protein detection in PFA-fixed paraffin-embedded SARS-CoV-2-infected COPD airway epithelium. STAR protocols. 2021;2(3):100663.

62. Coultas JA, Smyth R, Openshaw PJ. Respiratory syncytial virus (RSV): a scourge from infancy to old age. Thorax. 2019;74(10):986–93. Epub 2019/08/07. doi: 10.1136/thoraxjnl-2018-212212. PubMed PMID: 31383776.

63. Walsh D, Naghavi MH. Exploitation of Cytoskeletal Networks during Early Viral Infection. Trends Microbiol. 2019;27(1):39–50. Epub 2018/07/24. doi: 10.1016/j.tim.2018.06.008. PubMed PMID: 30033343; PubMed Central PMCID: PMCPMC6309480.

64. Chan KMC, Son S, Schmid EM, Fletcher DA. A viral fusogen hijacks the actin cytoskeleton to drive cell-cell fusion. Elife. 2020;9. Epub 2020/05/23. doi: 10.7554/eLife.51358. PubMed PMID: 32441254; PubMed Central PMCID: PMCPMC7244324.

65. Taylor MP, Koyuncu OO, Enquist LW. Subversion of the actin cytoskeleton during viral infection. Nat Rev Microbiol. 2011;9(6):427–39. Epub 2011/04/28. doi: 10.1038/nrmicro2574. PubMed PMID: 21522191; PubMed Central PMCID: PMCPMC3229036.

66. Matthews JD, Morgan R, Sleigher C, Frey TK. Do viruses require the cytoskeleton? Virol J. 2013;10:121. Epub 2013/04/20. doi: 10.1186/1743-422X-10-121. PubMed PMID: 23597412; PubMed Central PMCID: PMCPMC3639929.

67. Krzyzaniak MA, Zumstein MT, Gerez JA, Picotti P, Helenius A. Host cell entry of respiratory syncytial virus involves macropinocytosis followed by proteolytic activation of the F protein. PLoS Pathog. 2013;9(4):e1003309. Epub 2013/04/18. doi: 10.1371/journal.ppat.1003309. PubMed PMID: 23593008; PubMed Central PMCID: PMCPMC3623752.

68. Kallewaard NL, Bowen AL, Crowe JE, Jr. Cooperativity of actin and microtubule elements during replication of respiratory syncytial virus. Virology. 2005;331(1):73–81. Epub 2004/12/08. doi: 10.1016/j.virol.2004.10.010. PubMed PMID: 15582654.

69. Van Brusselen D, De Troeyer K, Ter Haar E, Vander Auwera A, Poschet K, Van Nuijs S, et al. Bronchiolitis in COVID-19 times: a nearly absent disease? Eur J Pediatr. 2021;180(6):1969–73. Epub 20210130. doi: 10.1007/s00431-021-03968-6. PubMed PMID: 33517482; PubMed Central PMCID: PMCPMC7847293.

70. Bara I, Ozier A, Tunon de Lara JM, Marthan R, Berger P. Pathophysiology of bronchial smooth muscle remodelling in asthma. Eur Respir J. 2010;36(5):1174–84. Epub 2010/11/03. doi: 10.1183/09031936.00019810. PubMed PMID: 21037369.

71. Florin TA, Plint AC, Zorc JJ. Viral bronchiolitis. Lancet. 2017;389(10065):211–24. Epub 2016/08/24. doi: 10.1016/S0140-6736(16)30951-5. PubMed PMID: 27549684; PubMed Central PMCID: PMCPMC6765220.

72. Li X, Sun S, Wu F, Shi T, Fan H, Li D. Study on JNK/AP-1 signaling pathway of airway mucus hypersecretion of severe pneumonia under RSV infection. Eur Rev Med Pharmacol Sci. 2016;20(5):853–7.

73. Lee J-W, Kim YI, Im C-N, Kim SW, Kim SJ, Min S, et al. Grape seed proanthocyanidin inhibits mucin synthesis and viral replication by suppression of AP-1 and NF-κB via p38 MAPKs/JNK signaling pathways in respiratory syncytial virus-infected A549 cells. Journal of Agricultural and Food Chemistry. 2017;65(22):4472–83.

74. Fonceca A, Flanagan B, Jeffers G, Smyth R, McNamara P. Mucin Expression in Airway Epithelial Cells from Children with Bronchiolitis, Healthy Children and RSV Infected Airway Epithelial Cell Cultures. D107 MECHANISMS OF MUCUS PRODUCTION DURING INFLAMMATION AND INFECTION: American Thoracic Society; 2009. p. A6311.

75. Johnson JE, Gonzales RA, Olson SJ, Wright PF, Graham BS. The histopathology of fatal untreated human respiratory syncytial virus infection. Mod Pathol. 2007;20(1):108–19. Epub 20061124. doi: 10.1038/modpathol.3800725. PubMed PMID: 17143259.

76. Groves H, Broadbent L, Guo-Parke H, Shields M, Power U. Influence of gestational and developmental age on human airway epithelial innate immune responses to Respiratory Syncytial Virus (RSV) in early life. Access Microbiology. 2019;1(1A):673.

77. Hillyer P, Shepard RE, Uehling M, Sheik F, Luongo C, Buchholz UJ, et al. Respiratory Syncytial Virus-Induced Host IFN Signaling Differs Between A549 and BEAS-2B Epithelial Cell Lines. Journal of Allergy and Clinical Immunology. 2015;135(2):AB9.

78. Yoboua F, Martel A, Duval A, Mukawera E, Grandvaux N. Respiratory syncytial virus-mediated NF-kappa B p65 phosphorylation at serine 536 is dependent on RIG-I, TRAF6, and IKK beta. J Virol. 2010;84(14):7267–77. Epub 2010/04/23. doi: 10.1128/JVI.00142-10. PubMed PMID: 20410276; PubMed Central PMCID: PMCPMC2898247.

79. Haeberle HA, Takizawa R, Casola A, Brasier AR, Dieterich HJ, Van Rooijen N, et al. Respiratory syncytial virus-induced activation of nuclear factor-kappaB in the lung involves alveolar macrophages and toll-like receptor 4-dependent pathways. J Infect Dis. 2002;186(9):1199–206. Epub 2002/10/29. doi: 10.1086/344644. PubMed PMID: 12402188.

80. McNamara PS, Flanagan BF, Hart CA, Smyth RL. Production of chemokines in the lungs of infants with severe respiratory syncytial virus bronchiolitis. J Infect Dis. 2005;191(8):1225–32. Epub 2005/03/19. doi: 10.1086/428855. PubMed PMID: 15776367.

81. Oshansky CM, Barber JP, Crabtree J, Tripp RA. Respiratory syncytial virus F and G proteins induce interleukin 1alpha, CC, and CXC chemokine responses by normal human bronchoepithelial cells. J Infect Dis. 2010;201(8):1201–7. doi: 10.1086/651431. PubMed PMID: 20205592; PubMed Central PMCID: PMCPMC2839062.

82. McNamara PS, Flanagan BF, Hart CA, Smyth RL. Production of chemokines in the lungs of infants with severe respiratory syncytial virus bronchiolitis. The Journal of infectious diseases. 2005;191(8):1225–32.

83. Machado D, Hoffmann J, Moroso M, Rosa-Calatrava M, Endtz H, Terrier O, et al. RSV infection in human macrophages promotes CXCL10/IP-10 expression during bacterial co-infection. International Journal of Molecular Sciences. 2017;18(12):2654.

84. Belperio JA, Keane MP, Burdick MD, Lynch JP, Xue YY, Li K, et al. Critical role for CXCR3 chemokine biology in the pathogenesis of bronchiolitis obliterans syndrome. The Journal of Immunology. 2002;169(2):1037–49.

85. Saetta M, Mariani M, Panina-Bordignon P, Turato G, Buonsanti C, Baraldo S, et al. Increased expression of the chemokine receptor CXCR3 and its ligand CXCL10 in peripheral airways of smokers with chronic obstructive pulmonary disease. American journal of respiratory and critical care medicine. 2002;165(10):1404–9.

86. Shahabuddin S, Ji R, Wang P, Brailoiu E, Dun N, Yang Y, et al. CXCR3 chemokine receptor-induced chemotaxis in human airway epithelial cells: role of p38 MAPK and PI3K signaling pathways. American Journal of Physiology-Cell Physiology. 2006;291(1):C34–C9.

87. Hardaker EL, Bacon AM, Carlson K, Roshak AK, Foley JJ, Schmidt DB, et al. Regulation of TNF-α and IFN-γ induced CXCL10 expression: participation of the airway smooth muscle in the pulmonary inflammatory response in chronic obstructive pulmonary disease. The FASEB journal. 2004;18(1):191–3.

88. Zeng Y, Wang R, Wang F, Zhang M, Zhang L, Zhu C, et al. Interaction of influenza A virus NS1 and cytoskeleton scaffolding protein α-actinin 4. Virus Genes. 2022;58(1):15–22.

89. Bamia A, Marcato V, Boissière M, Mansuroglu Z, Tamietti C, Romani M, et al. The NSs protein encoded by the virulent strain of Rift Valley fever virus targets the expression of Abl2 and the actin cytoskeleton of the host, affecting cell mobility, cell shape, and cell-cell adhesion. Journal of virology. 2020;95(1):e01768–20.

90. Armer H, Moffat K, Wileman T, Belsham GJ, Jackson T, Duprex WP, et al. Foot-and-mouth disease virus, but not bovine enterovirus, targets the host cell cytoskeleton via the nonstructural protein 3Cpro. Journal of virology. 2008;82(21):10556–66.

91. Liu H, Liu S, Liu Z, Gao X, Xu L, Huang M, et al. Dabie bandavirus Nonstructural Protein Interacts with Actin to Induce F-Actin Rearrangement and Inhibit Viral Adsorption and Entry. Journal of Virology. 2022:e00788–22.

92. Furnon W, Fender P, Confort M-P, Desloire S, Nangola S, Kitidee K, et al. Remodeling of the Actin network associated with the non-structural protein 1 (NS1) of West Nile virus and formation of NS1-containing tunneling nanotubes. Viruses. 2019;11(10):901.

93. Skoble J, Portnoy DA, Welch MD. Three regions within ActA promote Arp2/3 complex-mediated actin nucleation and Listeria monocytogenes motility. J Cell Biol. 2000;150(3):527–38. doi: 10.1083/jcb.150.3.527. PubMed PMID: 10931865; PubMed Central PMCID: PMCPMC2175181.

94. Dogan M, Han Y-S, Delmotte P, Sieck GC. TNFα enhances force generation in airway smooth muscle. American Journal of Physiology-Lung Cellular and Molecular Physiology. 2017;312(6):L994–L1002.

95. Shahriari S, Wei KJ, Ghildyal R. Respiratory Syncytial Virus Matrix (M) Protein Interacts with Actin In Vitro and in Cell Culture. Viruses. 2018;10(10). Epub 2018/10/03. doi: 10.3390/v10100535. PubMed PMID: 30274351; PubMed Central PMCID: PMCPMC6213044.

96. Le Nouen C, Munir S, Losq S, Winter CC, McCarty T, Stephany DA, et al. Infection and maturation of monocyte-derived human dendritic cells by human respiratory syncytial virus, human metapneumovirus, and human parainfluenza virus type 3. Virology. 2009;385(1):169–82. Epub 2009/01/09. doi: 10.1016/j.virol.2008.11.043. PubMed PMID: 19128816; PubMed Central PMCID: PMCPMC2668876.

97. Osan JK, DeMontigny BA, Mehedi M. Immunohistochemistry for protein detection in PFA-fixed paraffin-embedded SARS-CoV-2-infected COPD airway epithelium. STAR Protoc. 2021;2(3):100663. Epub 2021/07/13. doi: 10.1016/j.xpro.2021.100663. PubMed PMID: 34250510; PubMed Central PMCID: PMCPMC8259228.

98. Andrews S. FastQC: a quality control for high throughput sequence data. 2010 [cited 2019 October 7].

99. Bolger AM, Lohse M, Usadel B. Trimmomatic: a flexible trimmer for Illumina sequence data. Bioinformatics. 2014;30(15):2114–20. Epub 2014/04/04. doi: 10.1093/bioinformatics/btu170. PubMed PMID: 24695404; PubMed Central PMCID: PMCPMC4103590.

100. Trapnell C, Williams BA, Pertea G, Mortazavi A, Kwan G, van Baren MJ, et al. Transcript assembly and quantification by RNA-Seq reveals unannotated transcripts and isoform switching during cell differentiation. Nat Biotechnol. 2010;28(5):511–5. Epub 2010/05/04. doi: 10.1038/nbt.1621. PubMed PMID: 20436464; PubMed Central PMCID: PMCPMC3146043.

101. Liao Y, Smyth GK, Shi W. featureCounts: an efficient general purpose program for assigning sequence reads to genomic features. Bioinformatics. 2014;30(7):923–30. Epub 2013/11/15. doi: 10.1093/bioinformatics/btt656. PubMed PMID: 24227677.

102. Love MI, Huber W, Anders S. Moderated estimation of fold change and dispersion for RNA-seq data with DESeq2. Genome Biol. 2014;15(12):550. Epub 2014/12/18. doi: 10.1186/s13059-014-0550-8. PubMed PMID: 25516281; PubMed Central PMCID: PMCPMC4302049.

103. Yu G, Wang LG, Han Y, He QY. clusterProfiler: an R package for comparing biological themes among gene clusters. OMICS. 2012;16(5):284–7. Epub 2012/03/30. doi: 10.1089/omi.2011.0118. PubMed PMID: 22455463; PubMed Central PMCID: PMCPMC3339379.

